# *Toxoplasma gondii* encodes an array of TBC-domain containing proteins including an essential regulator that targets Rab2 in the secretory pathway

**DOI:** 10.1101/2023.05.28.542599

**Authors:** Justin J. Quan, Lachezar A. Nikolov, Jihui Sha, James A. Wohlschlegel, Peter J. Bradley

## Abstract

*Toxoplasma gondii* resides in its intracellular niche by employing a series of specialized secretory organelles that play roles in invasion, host-cell manipulation and parasite replication. Rab GTPases are major regulators of the parasite’s secretory traffic that function as nucleotide dependent molecular switches to control vesicle trafficking. While many of the Rab proteins have been characterized in *T. gondii*, precisely how these Rabs are regulated remains poorly understood. To better understand the parasite’s secretory traffic, we investigated the entire family of Tre2–Bub2–Cdc16 (TBC)-domain containing proteins, which are known to be involved in vesicle fusion and secretory protein trafficking. We first determined the localization of all 18 TBC-domain containing proteins to discrete regions of the secretory pathway or other vesicles in the parasite. We then use an auxin-inducible degron approach to demonstrate that the protozoan-specific TgTBC9 protein that localizes to the ER is essential for parasite survival. Knockdown of TgTBC9 results in parasite growth arrest and affects the organization of the ER and Golgi apparatus. We show that the conserved dual-finger active site in the TBC-domain of the protein is critical for its GTPase-activating protein (GAP) function and that the *P. falciparum* orthologue of TgTBC9 can rescue the lethal knockdown. We additionally show by immunoprecipitation and yeast two hybrid analyses that TgTBC9 directly binds Rab2, indicating that this TBC-Rab pair controls ER to Golgi traffic in the parasite. Together, these studies identify the first essential TBC protein described in any protozoan, provide new insight into intracellular vesicle trafficking in *T. gondii*, and reveal promising targets for the design of novel therapeutics that can specifically target apicomplexan parasites.

## Introduction

*Toxoplasma gondii* is an obligate intracellular parasite in the phylum Apicomplexa that causes the disease toxoplasmosis and infects all mammals including approximately one-third of the human population.^1,2^ Although most human infections remain asymptomatic, immunocompromised individuals or congenitally infected neonates are vulnerable to more severe symptoms such as blurred vision, seizures, coordination issues, lung problems, or fatality if left untreated.^3^ *T. gondii* is the most widespread apicomplexan and serves as a model system for studying other apicomplexan parasites, such as *Cryptosporidium spp.*, which causes diarrheal disease, and *Plasmodium falciparum*, which causes malaria.^4,5^ In addition to the standard eukaryotic organelles, apicomplexans share a number of unique organelles such as the inner membrane complex (IMC), micronemes, rhoptries, apicoplast, and dense granules that facilitate host cell invasion and intracellular survivial.^6,7^ As these unique organelles often contain parasite-specific proteins, they are considered excellent targets for therapeutic intervention.

One interesting aspect of *T. gondii* is its highly polarized secretory pathway, which delivers secretory proteins to organelles at the apical end of the parasite.^8,9^ As in other eukaryotic cells, secretory traffic is initiated by cotranslational translocation of protein cargo into the endoplasmic reticulum (ER). In *T. gondii*, this traffic then passes through a single Golgi stack which is positioned just anterior to the parasite’s nucleus.^7^ After exiting the Golgi, vesicle-based trafficking allows the protein cargo to be directed through post-Golgi compartments to the secretory micronemes, rhoptries, and dense granules, which discharge their contents sequentially for motility, host cell invasion and modification of the nascent parasitophorous vacuole.^7,8^ In addition to the secretory organelles, protein cargo from the secretory pathway is also targeted to the IMC, apicoplast, plant-like vacuolar compartment (PLVAC), endosome-like compartment (ELC), or parasite’s surface.^7,10–12^ Specific signals have been identified for targeting to most of these compartments, as has some of the vesicle trafficking machinery.^10,13^ However, a detailed mechanistic understanding of vesicle sorting as well as the identification of many components of the trafficking machinery has yet to be completed.

Similar to other eukaryotic cells, one of the major regulators of *Toxoplasma* secretory traffic is the Rab family of small GTPases.^14^ Rab proteins are tethered to membranes by C-terminal prenylation and function as molecular switches that cycle between an inactive GDP-bound state and an active GTP-bound state to regulate secretory and vesicular traffic.^15^ Rabs are regulated by guanine nucleotide exchange factors (GEFs) that promote the exchange of GDP for GTP as well as GTPase activating proteins (GAPs), which hydrolyze GTP to GDP.^16^ Twelve Rab proteins have been identified in *T. gondii*, which appear to be conserved across most of the eukarya.^17,18^ The proteins localize to various secretory or vesicular compartments such as the ER/Golgi, Golgi, post Golgi compartments, or cytoplasmic vesicles.^17^ Functional assessment of the Rabs, by dominant negative or overexpression screening, has confirmed their importance in protein trafficking and parasite fitness.^17^ For example, overexpression of Rab2, Rab4, Rab5A, Rab5B and Rab5C dramatically affect parasite growth and overexpression of Rab5A and Rab5C results in mistargeting of rhoptry and microneme cargo into the vacuole.^17,19,20^ Rab11A has been shown to regulate dense granule secretion and Rab11B functions in IMC biogenesis.^21,22^ In addition, some of the GEFs that activate Rabs have been described including a Vps9 and a Vps11 domain containing protein, although their target Rabs have not been identified.^23,24^

While several Rab GEFs have been studied in *T. gondii*, the GAP proteins responsible for inactivating Rabs have not been explored. Rab GAPs are typically characterized by the presence of a TBC (Tre-2/Bub2/Cdc16) domain and act by increasing the intrinsic GTPase activity of the Rab protein, resulting in a conversion from the GTP to GDP bound form.^25^ TBC domain-containing proteins (hereafter termed TBC proteins) often contain other functional domains that presumably contribute to their function in various locations within the cell.^25^ The TBC family is poorly studied in most systems as there are typically many family members that often have similar localizations, suggesting functional redundancy (e.g. there are 41 TBC proteins in humans).^25,26^ In addition, there are a small number of other proteins that lack a TBC domain but can still function as Rab GAPs.^16,27^ The TBC proteins in most parasites remain largely unstudied, and none have been localized or functionally assessed in *T. gondii*. In addition, phylogenetic analyses suggest that the most recent common ancestor of the eukaryotes contained several TBC proteins, which gave rise to a number of TBC clades in different eukaryotic lineages.^14^ The relationships among clades are poorly resolved likely due to their ancient divergences, which complicates studies of TBC family proteins as clear orthologues are frequently difficult to determine.

Given the gap in our understanding of vesicular transport regulators and GAPs, we performed a systematic analysis of TBC proteins in *T. gondii*. In this study, we identify 18 TBC proteins and localize them to discrete compartments of the secretory pathway or other vesicles in the parasite. We then focus on the ER-localized TgTBC9 and demonstrate that conditional knockdown of this family member results in a lethal growth arrest and disruption of the ER and Golgi. We also show by mutagenesis that the GAP activity of TgTBC9 is critical for function and that the *P. falciparum* orthologue of TgTBC9 can rescue the knockdown. We also show that TgTBC9 directly binds to the ER-Golgi localized Rab2 protein, indicating this is the major target of TgTBC9.^17^ This work substantially builds on our understanding of vesicular protein targeting in *T. gondii* and identifies a TBC protein that is essential for cellular survival. TgTBC9 is the first essential TBC protein identified in any protozoan, thus revealing a critical player in secretory traffic and a new potential drug target for these organisms.

## Results

### *T. gondii* encodes 18 TBC proteins that are mostly unique to the Apicomplexa

To identify the repertoire of TBC proteins in *T. gondii*, we searched for proteins that contain a TBC domain as predicted by NCBI, InterPro, and SMART conserved domain searches^28–30^. This reveals a total of 18 TBC proteins which range from a molecular weight of 35 kDa to 426 kDa (Figure 1; Table 1). Nine of these are over 200 kDa in size, which is considerably larger than most TBC proteins (e.g. human TBC proteins range from 34.97 kDa (TBC1D21) to 158.69 kDa (UPS6)). Similar to that seen in other systems, the TBC domain is situated at various positions in the proteins, and they frequently contain additional identifiable protein motifs such as coiled-coil (CC) domains, transmembrane domains (TM), kinase domains, or Sec7 catalytic domains (Figure 1). The Sec7 domain is a well characterized motif belonging to the GEF family, which are known to activate Rabs.^31^ As TgTBC15 and TgTBC16 contain both a Sec7 domain and TBC domain, they harbor both positive and negative regulators of Rab proteins (Figure 1). Members of this novel subfamily of the TBC proteins that contain both TBC and Sec7 domains are often called TBC-Sec7 (TBS) proteins and are only found in the protists.^32^

**Figure 1.**
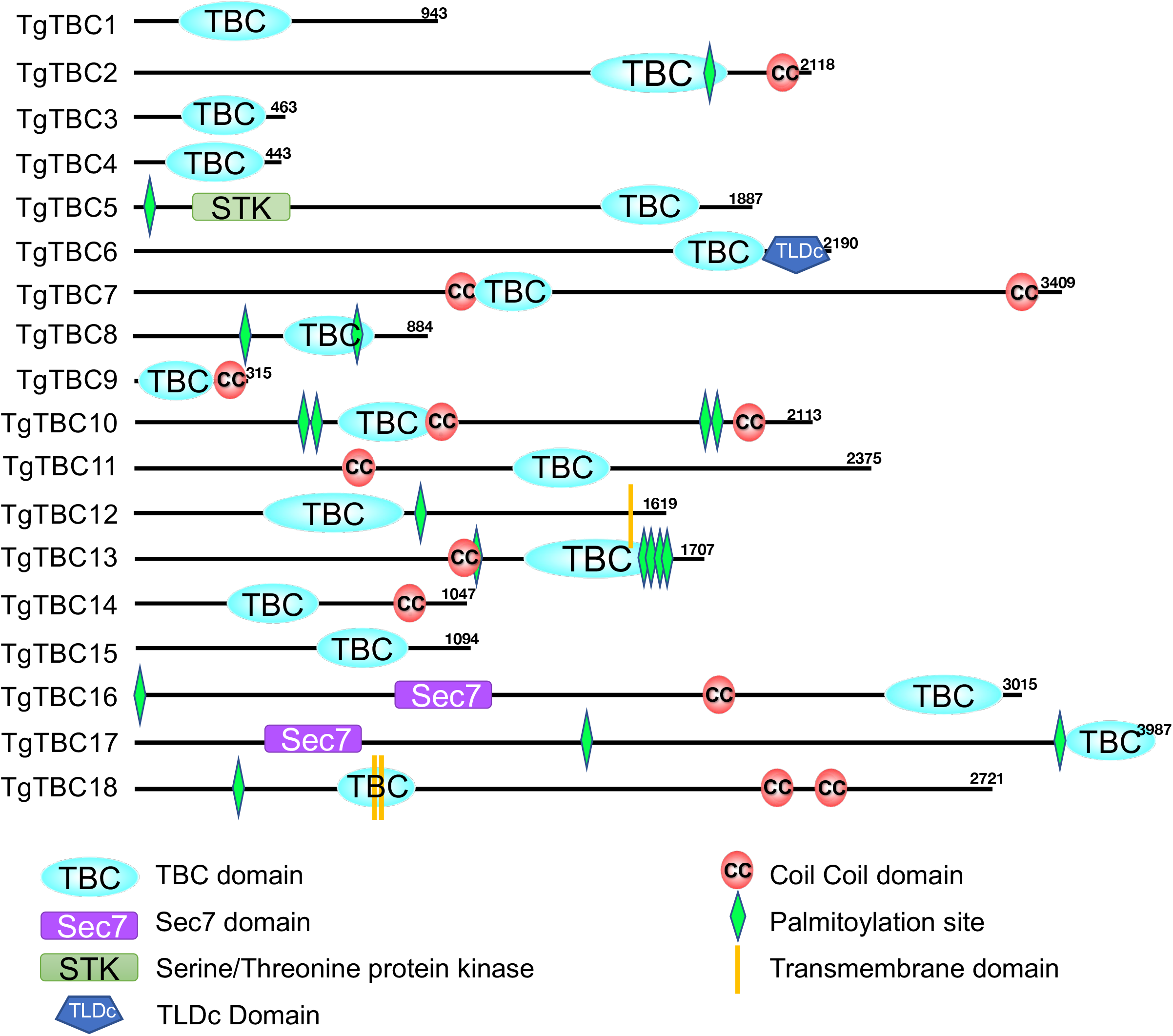
*Toxoplasma gondii* contains 18 TBC proteins. Diagram showing the domains present in the *Toxoplasma* TBC proteins. The approximate position of the TBC and other domains as predicted by SMART (Simple Modular Architecture Research Tool), PFAM, and NCBI conserved protein domain search tools are shown.^28–30^ TBC, Tre2–Bub2–Cdc16 (TBC)-domain containing proteins; Sec7, Sec7-domain-containing; STK, Serine/Threonine protein kinase; TLDc, <u>T</u>BC, <u>L</u>ysM, <u>D</u>omain <u>C</u>atalytic domain; CC, coil-coil domain; TM, transmembrane.

**Table 1.** Summary of 18 TBC-domain containing proteins in *Toxoplasma gondii*.

| Gene Name | ToxoDB-ID | Protein Size (kDa)* | Phenotype Score** | Localization (Tachyzoite) | IxxDxxR Predicted Active Site | YxQ Predicted Active Site |
| --- | --- | --- | --- | --- | --- | --- |
| TgTBC1 | TgGT1_275350 | 102 | -0.58 | Golgi | Yes | Yes |
| TgTBC2 | TgGT1_274130 | 226 | -1.84 | Golgi | No | Yes |
| TgTBC3 | TgGT1_218870 | 52 | -0.74 | Golgi | Yes | Yes |
| TgTBC4 | TgGT1_285730 | 49 | -0.36 | Not expressed | Yes | Yes |
| TgTBC5 | TgGT1_250680 | 207 | -0.06 | Cytoplasmic Vesicles | No | No |
| TgTBC6 | TgGT1_237280 | 233 | -0.16 | ER | No | No |
| TgTBC7 | TgGT1_216430 | 358 | 0.09 | Not expressed | Yes | Yes |
| TgTBC8 | TgGT1_261200 | 99 | -1 | IMC | Yes | Yes |
| TgTBC9 | TgGT1_226550 | 35 | -4.82 | ER | Yes | Yes |
| TgTBC10 | TgGT1_203910 | 228 | 0.53 | Cytosolic | Yes | Yes |
| TgTBC11 | TgGT1_239830 | 257 | -2.83 | Daughter | No | Yes |
| TgTBC12 | TgGT1_223640 | 178 | -0.6 | Cytoplasmic Vesicles | Yes | Yes |
| TgTBC13 | TgGT1_221710 | 182 | -2.04 | Cytoplasmic Vesicles | No | Yes |
| TgTBC14 | TgGT1_289820 | 114 | -1.14 | ER | Yes | Yes |
| TgTBC15 | TgGT1_226850 | 117 | 0.54 | Daughter | Yes | Yes |
| TgTBC16 | TgGT1_312300 | 324 | -1.19 | Cytoplasmic Vesicles | No | No |
| TgTBC17 | TgGT1_266830 | 426 | 0.3 | Cytoplasmic Vesicles | No | Yes |
| TgTBC18 | TgGT1_213325 | 293 | -2.86 | Golgi | Yes | Yes |

\*Predicted molecular weight of open reading frame without post-translational modification and epitope tagging.
\*\*Genome-wide CRISPR screen (GWCS) (27). A negative score indicates impaired growth.

Because standard phylogenetic analyses of TBC proteins have been shown to provide limited information, we used the OrthoMCL database to identify orthologues that could be clearly determined (Figure S1A; Table S1).^33^ We also used BLAST analysis to identify several likely orthologues that were not predicted by OrthoMCL.^34^ This showed that TgTBC1-8 have orthologues that are present in a broad range of eukaryotic lineages, suggesting these proteins may carry out functions in common to their eukaryotic counterparts. In contrast, TgTBC9-18 appear to be restricted to protozoans, with TgTBC13-18 being unique to *Toxoplasma* and its closest relatives (e.g. *Neospora caninum*, but not other coccidians or apicomplexans). We also found that ten of the *T. gondii* TBC proteins contain a recognizable dual-finger active site consisting of an “arginine finger” (IxxDxxR) and a “glutamine finger” (YxQ), suggesting TBC domain catalytic activity (Figure S1B; Table 1).^35^

### *Toxoplasma* TBC proteins localize to the secretory pathway and cytoplasmic vesicles

To determine the localization of TBC proteins in *T. gondii*, we endogenously tagged the C-terminus of each TBC protein with an epitope tag in tachyzoites (Figure 2A). Proteins with higher expression levels were tagged with a 3xHA, 3xTy, or 3xMyc tag, while those that were likely to have low levels of expression were tagged with either a spaghetti monster HA (smHA) or spaghetti monster OLLAS (smOLLAS) epitope tag.^36,37^ Using this approach, we found that TgTBC1, 2, 3 and 18 localizes to a bar-like shape that is positioned anterior to the nucleus, which suggested localization to the Golgi or Golgi-adjacent compartments. We co-stained these four proteins with the cis-Golgi marker GRASP55 and observed varying degrees of overlap (Figure 2B).^38^ TgTBC1 and 2 colocalize best with GRASP55, while TgTBC3 and 18 appear slightly more posterior, indicating that these localize to trans-Golgi or post-Golgi compartments.^38^ Eight of the TBC proteins (TgTBC5, 6, 9, 12, 13, 14, 16, 17) showed a nuclear-excluded, reticular or spotted cytoplasmic staining that suggested the endoplasmic reticulum (ER) or cytoplasmic vesicles. We co-stained these with the ER marker SERCA and while it is difficult to make definitive localizations, TgTBC6, 9, and 14 colocalized best and were considered most likely ER (Figure 2C).^39^ The remaining five were denoted as cytoplasmic vesicles (Figure 2D), some of which likely represent different levels of intersection of the endocytic and exocytic pathways which converge downstream of the Golgi, although localization to other types of cytoplasmic vesicles is also possible. TgTBC10 localizes to spots in the cytoplasm, but it is not nuclear-excluded as assessed by SERCA co-staining (Figure 2E).

**Figure 2.**
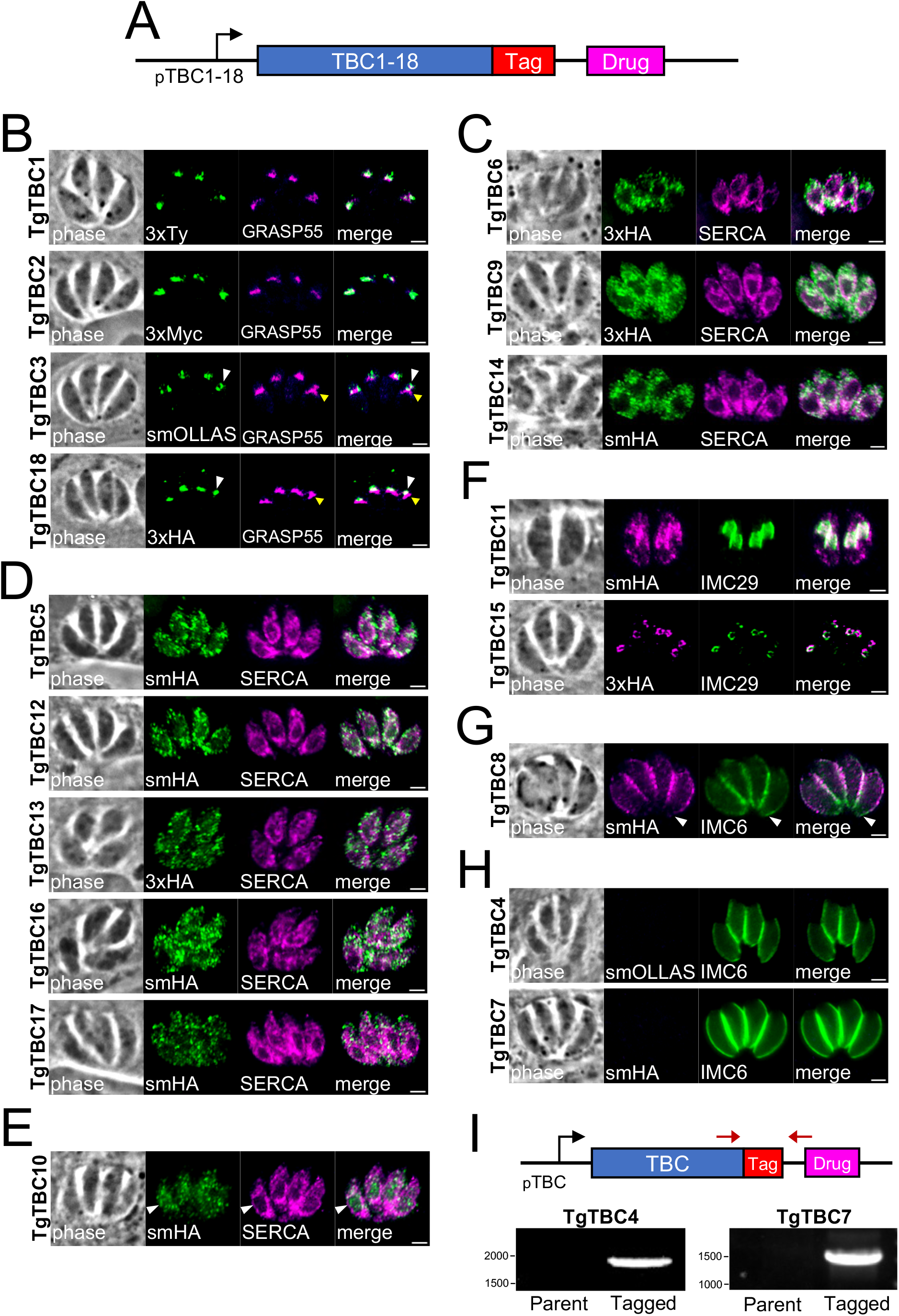
*T. gondii* TBC proteins localize to discrete regions of the secretory pathway and cytoplasmic vesicles. IFAs of endogenously epitope tagged TgTBC1-18 parasites. A) Diagram of TgTBC1-18 showing the epitope tag and selectable marker. B) IFA of endogenously tagged TgTBC1, 2, 3, and 18 stained with antibodies against epitope tags and colocalized with GRASP55. The yellow and white arrowheads in the bottom two panels denote subtle differences between the TBC protein and GRASP55. Green = endogenously tagged TBC proteins, Magenta = GRASP55-mCherry. C) IFA of TgTBC6, 9 and 14 stained with anti-HA and colocalized with SERCA. Green = rabbit anti-HA, Magenta = mouse anti-SERCA. D) IFA of TgTBC 5, 12, 13, 16, and 17 shown partially colocalizing with SERCA. Green = rabbit anti-HA, Magenta = mouse anti-SERCA. E) IFA of TgTBC10 shows cytoplasmic and nuclear staining with partial colocalization with SERCA. The white arrowheads show that TgTBC10 is not nuclear excluded. Green = rabbit anti-HA, Magenta = mouse anti-SERCA. F) IFA of TgTBC11 and TgTBC15 colocalizing with endogenously tagged IMC29^3xV5^ in daughter buds. Magenta = mouse anti-HA, Green = rabbit anti-V5. G) IFA of TgTBC8 shows staining central portion of the maternal IMC as accessed by IMC6 staining. The white arrowheads indicate basal portions of the parasite where TgTBC8 is absent. Magenta = mouse anti-HA, Green = rabbit anti-IMC6. H) IFA of TgTBC4 and TgTBC7 with no detectable smHA staining. Magenta = mouse anti-HA, Green = rabbit anti-IMC6. Scale bars for all images, 2 μm. I) Diagram and PCR of endogenous TgTBC4 and TgTBC7 tagged parasites. Primers labeled as red arrows were used to test gDNA from parental and tagged strains.

In addition, three of the TBC proteins were found to localize to the parasite-specific IMC. Two of these localized to the daughter buds of IMC during endodyogeny, as assessed by colocalization with the early daughter bud marker IMC29 (Figure 2F).^40^ In contrast, TgTBC8 appears to localize to the maternal IMC, as it is peripheral but not present in the apical cap or basal portion of the organelle (Figure 2G). Finally, in agreement with the low expression levels suggested by ToxoDB, TgTBC4 and 7 could not be detected even though integration of the tag was confirmed by PCR, indicating that these family members are not significantly expressed in the parasite’s tachyzoite stage (Figure 2H and 2I).^41,42^

### TgTBC9 is essential for parasite survival and organization of the ER and Golgi

We focused on TgTBC9 because it was assigned the lowest phenotype score (−4.82) of the TBC proteins in the *Toxoplasma* genome-wide CRIPSR screen (GWCS), suggesting that the protein is either important for parasite fitness or essential (Table 1).^43^ Phylogenetic analysis based on the conserved TBC domain of homologs from apicomplexans, amebozoans and trypanosomatids demonstrates that TgTBC9 is a member of a protozoan-restricted clade of TBC proteins previously denoted as TBC-RootA (Figure S1A).^25^ Neighbor joining analysis shows that TgTBC9 forms a well-supported clade with TBC-RootA homologs from other apicomplexans (*P. falciparum, T. parva*, *C. parvum*) (Figure 3A). To determine the function of TgTBC9, we used a conditional knockdown approach by generating parasites with TgTBC9 endogenously tagged with the mIAA7 auxin-inducible degron (AID) fused to 3xHA (TgTBC9^AID-3xHA^) (Figure 3B). The mIAA7 degron tag was chosen as it is known to be less susceptible to basal degradation in the absence of auxin (3-indoleacetic acid, IAA).^44^ The TgTBC9^AID-3xHA^ fusion protein localized to an ER-like pattern similar to that seen for the 3xHA tagged version (Figure 2C and 3C). To assess the effects of the TgTBC9 knockdown on parasite growth, intracellular parasites were treated with and without IAA for 4 hours, syringe lysed, and then allowed to infect confluent monolayers for 24 hours with and without IAA. Using this approach, we observed that TgTBC9 protein levels decreased to an undetectable level by IFA and western blot, demonstrating an effective depletion of the protein (Figure 3D and 3E). Replication of the knockdown parasites appears to cease, often arresting at two parasites per vacuole, although the overall morphology of the arrested parasites generally remains unaffected as assessed by IMC6 staining (Figure 3D). To evaluate the knockdown at a longer time point, we conducted plaque assays and found that TgTBC9^AID-3xHA^ parasites were unable to form plaques upon IAA treatment, demonstrating that TgTBC9 is essential for parasite survival (Figure 3F and 3G).

**Figure 3.**
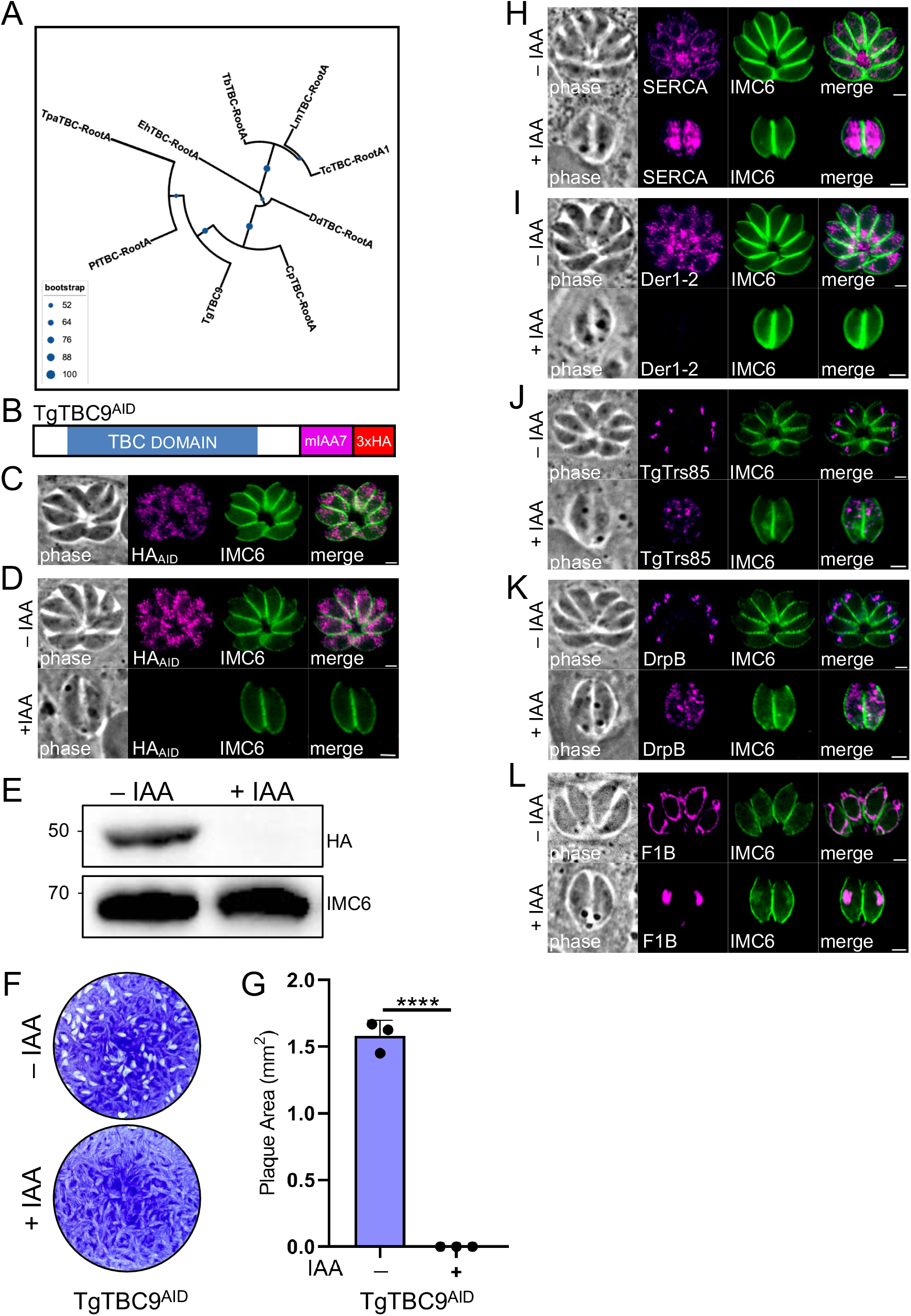
TgTBC9 is essential for parasite survival and organization of the ER and Golgi. A) Maximum likelihood tree based on the TBC domains of TgTBC9 and its orthologs from *Plasmodium falciparum* (Pf), *Trypanosoma brucei* (Tb), *Entamoeba histolytica* (Eh), *Cryptosporidium parvum* (Cp), *Theileria parva* (Tp), *Trypanosoma cruzi* (Tc), *Leishmania major* (Lm), *Dictyostelium discoideum* (Dd). Blue spheres denote level of support (1000 bootstrap replicates). B) Diagram of TgTBC9^AID^ showing its TBC domain and a mIAA7-3xHA degron tag added to the C-terminus of the protein. C) IFA of TgTBC9^AID^ localizing to an ER-like pattern. Magenta, mouse anti-HA; green, rabbit anti-IMC6. D) IFA of TgTBC9^AID^ without (−) or with (+) IAA for 24 h (following a 4 h pretreatment ±IAA) showing that TgTBC9^AID^ is efficiently degraded, resulting in parasite growth arrest. Magenta, mouse anti-HA; green, rabbit anti-IMC6. E) Western blot analysis of showing TgTBC9^AID^ is efficiently degraded upon IAA treatment. IMC6 is used as a load control. F) Plaque assays showing that TgTBC9-depleted parasites are unable to form plaques. G) Quantification of plaque assays at day 7 showing no plaque formation by TgTBC9^AID^ parasites +IAA. H) IFA of TgTBC9^AID^ parasites grown in ±IAA as described in D showing affected ER morphology. Magenta, mouse anti-SERCA; green, rabbit anti-IMC6. I) TgTBC9^AID-3xTy^ parasites grown in ±IAA as described in D but with staining for Der1-2 using an endogenously tagged Der1-2^3xHA^ strain. Magenta, mouse anti-HA; green, rabbit anti-IMC6. J) TgTBC9^AID^ parasites grown in ±IAA as described in D but with staining for Golgi using endogenously tagged TgTrs85^3xV5^ strain. Magenta, mouse anti-V5; green, rabbit anti-IMC6. K) TgTBC9^AID^ parasites grown in ±IAA as described in D but with staining for DrpB. Magenta, rat anti-DrpB; green, rabbit anti-IMC6. L) TgTBC9^AID^ parasites grown in ±IAA as described in D but with staining for mitochondrion using anti-F1β ATPase. Magenta, mouse anti-F1β ATPase; green, rabbit anti-IMC6. (****, *P <* 0.0001). Scale bars for all images, 2 μm.

Since the TgTBC9 knockdown parasites arrest, we examined a series of organelles in the knockdown strain, including the ER, Golgi, post-Golgi compartment labeled by DrpB, PLVAC, ELC, micronemes, rhoptries, dense granules, IMC, centrosome, apicoplast, and mitochondria. Consistent with the degron tagged protein localizing to the ER, we found TgTBC9 knockdown causes the ER to lose its reticular pattern and become more dense in the cytoplasm as assessed by staining with the ER marker SERCA (Figure 3H).^39^ We also examined the ER marker Der1-2 and found that it was largely lost upon knockdown, again indicating that the ER is affected upon the loss of TgTBC9 (Figure 3I).^45^ We then examined the *medial*-Golgi protein TgTrs85 and the post-Golgi compartment labeled by DrpB.^11,46^ Both Golgi associated structures were also disrupted with diffuse spots in the cytoplasm instead of a more discrete bar-like structure anterior to the nucleus (Figure 3J and 3K). Interestingly, the rhoptries, micronemes, dense granules, PLVAC, ELC, apicoplast, centrosome, and IMC all appear to remain unaffected (Figure S2). We did observe that TgTBC9 knockdown affected the morphology of the mitochondria, which collapsed from its typical lasso-like shape into a single rounded organelle (Figure 3L). This may suggest the parasite is losing viability, but a similar collapse of the mitochondria is seen when the mitochondria-IMC membrane tether is lost without leading to parasite death.^47,48^ Taken together, these data indicate that TgTBC9 plays a critical role in the parasite’s secretory protein trafficking and is essential for parasite growth.

### TBC dual-finger active sites are required for TgTBC9 GAP activity

TgTBC9 is one of the TBC proteins that contains a conserved dual-finger active site with the IxxDxxR and YxQ motifs (Fig S1; Table 1). To investigate whether the active site residues are required for TgTBC9’s GAP activity, we generated complementation constructs with the full length wild-type gene (wt), a mutant of the IxxDxxR motif (R74A), or a mutant of the YxQ motif (Q101A) (Figure 4A), which have previously been shown to disrupt function in other systems.^35,49^ Complementation with the wild-type gene (TgTBC9^wt^) in TgTBC9^AID-3xHA^ parasites restored the parasite’s ability to form plaques in the presence of IAA (Figure 4B and C). In contrast, complementation with either the TgTBC9^R74A^ or TgTBC9^Q101A^ mutant completely failed to rescue the parasite’s ability to form plaques upon knockdown of endogenous TgTBC9 (Figure 4D). Failure of the mutants to complement the knockdown was not due to expression levels, as similar levels of the TgTBC9^wt^ and the TgTBC9^R74A^ and TgTBC9^Q101A^ point mutants were confirmed by western blot analysis (Figure 4E). Quantification of these results confirmed that mutation of the TgTBC9 active sites completely disrupts the protein’s ability to rescue the lethal knockdown (Figure 4F).

**Figure 4.**
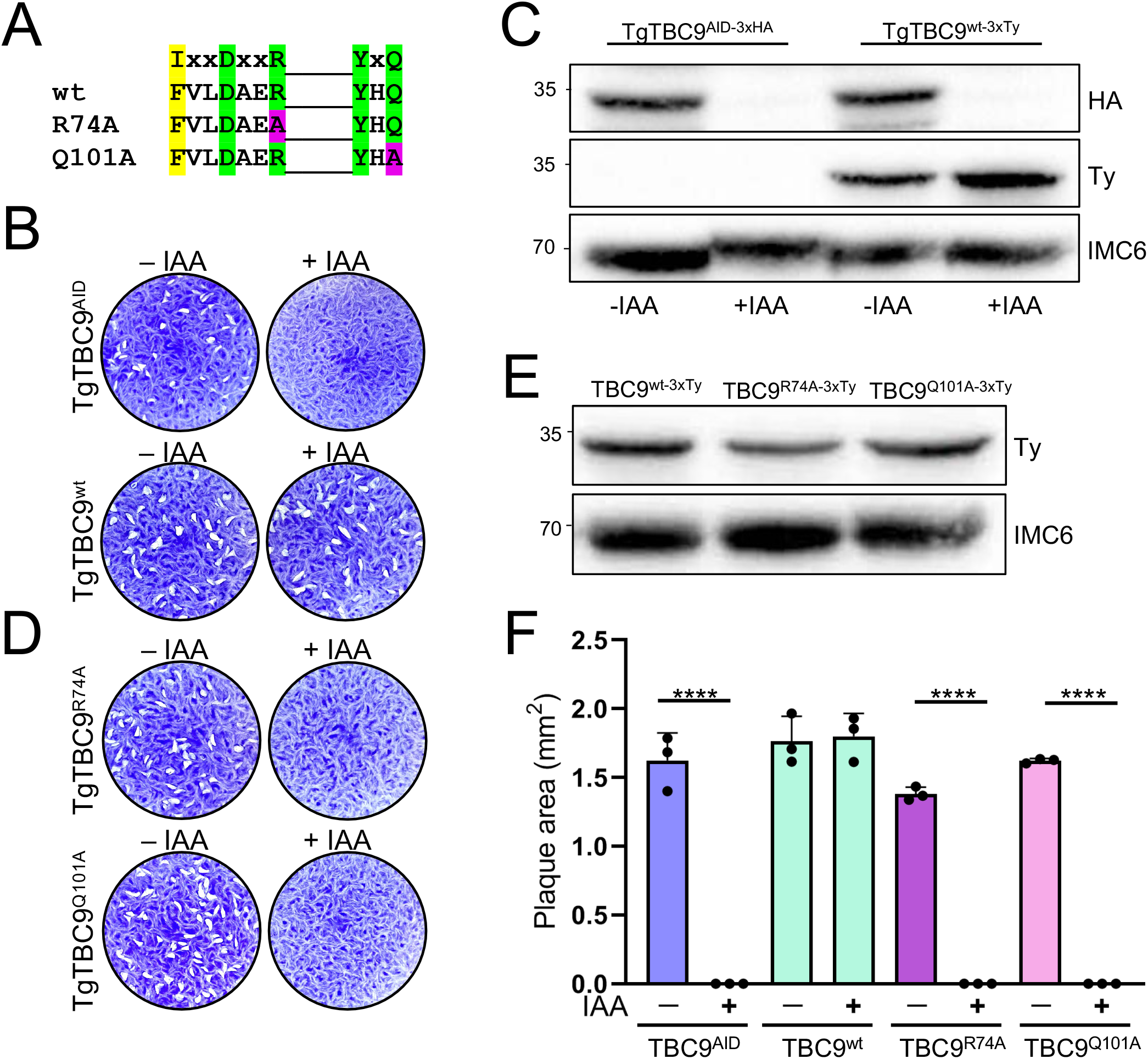
TBC dual-finger active sites are required for TgTBC9 GAP activity. A) Diagram of the dual-finger consensus and highlighting which TgTBC9 residues in the arginine finger and glutamine finger motifs were mutated to alanine. Green boxes depict strictly conserved residues; yellow boxes depict semi-conserved residues; magenta boxes depict residues that were mutated to alanine. B) Plaque assays showing that complementation with full-length TgTBC9 restores ability to form plaques upon depletion of endogenous TgTBC9. C) Western blot analysis showing knockdown of endogenous TgTBC9 and complementation with a Ty-tagged TgTBC9^wt^ copy targeted to the UPRT locus. IMC6 is used as a load control. D) Plaque assays showing complementation with the TgTBC9^R74A^ or TgTBC9^Q101A^ mutants are unable to rescue the TgTBC9^AID^ knockdown. E) Western blot analysis showing that the TgTBC9^wt^ and TgTBC9^R74A^ and TgTBC9^Q101A^ mutant parasites have similar levels of expression. IMC6 is used as a load control. F) Quantification of plaque assays at day 7 showing rescue with TgTBC9^wt^ but no plaque formation by TgTBC9^AID^, TgTBC9^R74A^, and TgTBC9^Q101A^ mutant parasites +IAA (****, *P* ≤ 0.0001).

### PfTBC9 partially rescues the lethal knockdown of TgTBC9 in *T. gondii*

As predicted by the OrthoMCL database, TgTBC9 has an ortholog in *Plasmodium falciparum* (PF3D7_0904000) that we denoted PfTBC9 (Figure S1A; Table S1). Surprisingly, the alignment between TgTBC9 and PfTBC9 reveals remarkable similarity with the exception of a 112 amino acid C-terminal extension in PfTBC9 (Figure 5A). Like its *Toxoplasma* counterpart, PfTBC9 is predicted to be essential and has the dual-finger active site.^50^ To test if PfTBC9 could rescue function, we cloned a codon-optimized PfTBC9 into our complementation vector and expressed it in TgTBC9^AID^ parasites. However, despite driving the gene from several promoters, we found that the construct appeared to start from an internal methionine and made a truncated protein that likely lacked the active site (not shown). To circumvent this problem, we constructed an N-terminal YFP construct for PfTBC9 and expressed it in the TgTBC9^AID^ strain (Figure 5B). The YFP tagged protein showed a similar ER-like pattern and was expressed at the correct size (Figure 5C-E). In this context, PfTBC9 was only partially able to rescue the lethal phenotype resulting in the formation of smaller plaques compared to complementation with the TgTBC9^wt^ gene, but no plaque efficiency defect was observed (Figure 5F-H). To determine if this partial rescue was a consequence of the YFP fusion, we similarly expressed TgTBC9^wt^ as a YFP fusion in the TgTBC9^AID^ strain and found that it also resulted in small plaques (Figure 5I-K). These results indicate that the N-terminal YFP fusion partially disrupts function and that PfTBC9 is able to complement the knockdown to a similar extent as TgTBC9.

**Figure 5.**
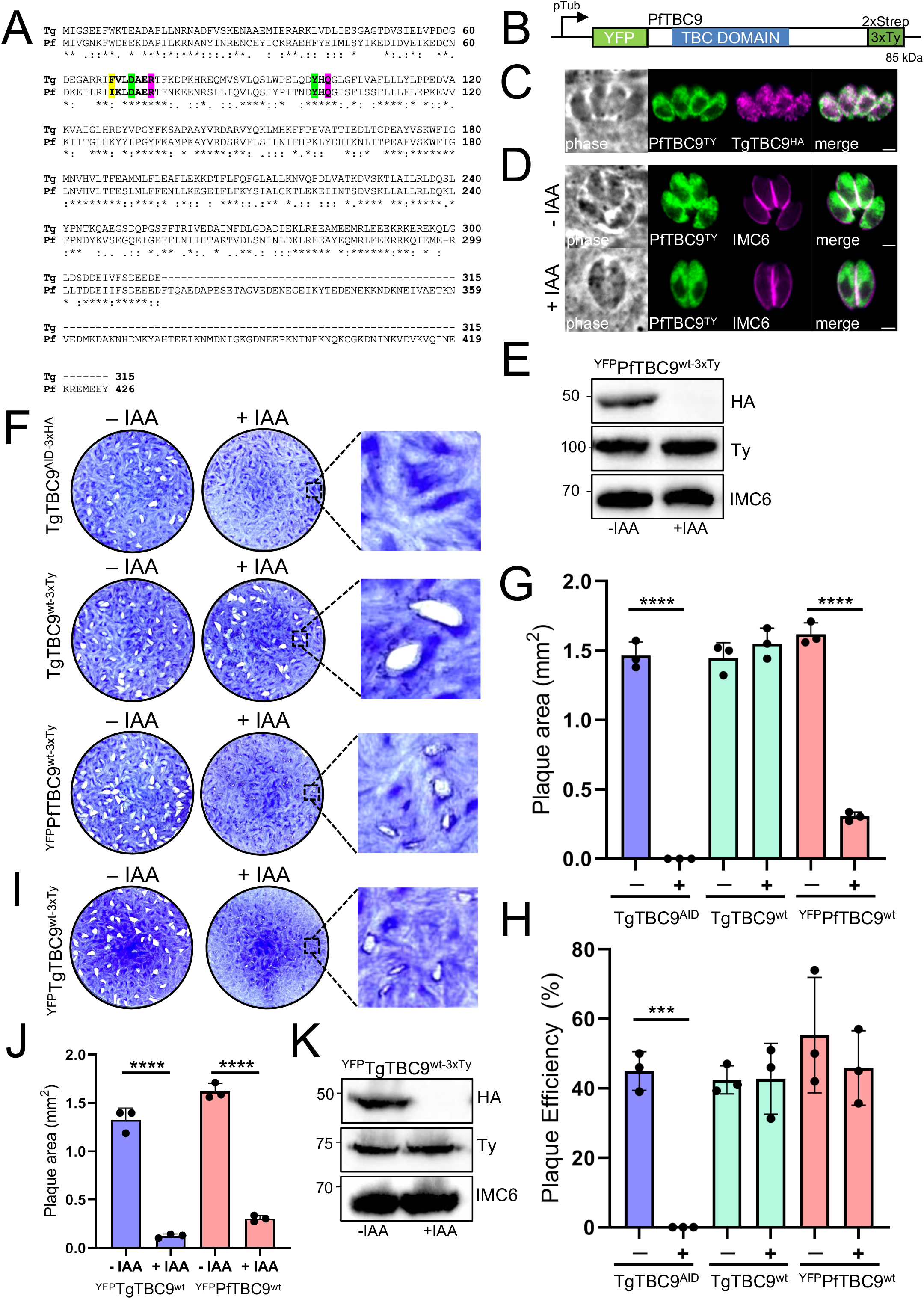
PfTBC9 partially rescues the lethal knockdown of TgTBC9. A) Full protein alignment of TgTBC9 (Tg) and PfTBC9 (Pf) using Clustal Omega.^70^ Bold residues highlighted in yellow depict semi-conserved residues; bold residues in green depict strictly conserved residues; bold residues in magenta indicate predicted R and Q catalytic sites. B) Diagram of ^YFP^PfTBC9^wt-3xTy^ expressed from the tubulin promoter with a N-terminal YFP and C-terminal 2xStrep-3xTy tag. C) IFA of ^YFP^PfTBC9^wt-3xTy^ colocalized with TgTBC9^AID-3xHA^ showing overlap in the ER. Green, mouse anti-Ty and GFP; magenta, rabbit anti-HA. Scale bar = 2 μm. D) IFA of PfTBC9^wt^ with (−) or without (+) IAA for 24 h showing that PfTBC9^wt^ is expressed and unaffected in ±IAA. Magenta, mouse anti-Ty and GFP; green, rabbit anti-HA. Scale bar = 2 μm. E) Western blot analysis of ^YFP^PfTBC9^wt-3xTy^ in the background of TgTBC9^AID^ tagged parasites. IMC6 is used as a load control. F) Plaque assays showing that PfTBC9^wt^ complemented parasites formed smaller plaques compared to the TgTBC9^wt^ complemented parasites. G) Quantification of plaque area at day 7 showing small plaque formation by ^YFP^PfTBC9^wt^ complemented parasites +IAA (****, *P <* 0.0001). H) Quantification of plaque efficiency at day 7 show no significance between ^YFP^PfTBC9^wt-3xTY^ +IAA from TgTBC9^wt^ complement groups (***, *P =* 0.0002). I) Plaque assays showing that ^YFP^TgTBC9^wt-3xTy^ complemented parasites +IAA similarly form small plaques. J) Quantification of plaque area at day 7 showing small plaque formation by ^YFP^TgTBC9^wt^ and ^YFP^PfTBC9^wt^ complemented parasites (****, *P <* 0.0001). K) Western blot analysis of ^YFP^TgTBC9^wt-3xTy^ in the background of TgTBC9^AID^ tagged parasites. IMC6 is used as a load control.

The *Trypanosoma brucei* orthologue of TgTBC9 is also a member of the RootA clade which we denote TbTBC9 (Tb427.10.7680) (Figure S1; Table S1 and S2).^25^ Interestingly, TbTBC9 has a 200 amino acid N-terminal extension and also contains the dual-finger active site (Figure S3A and B). However, TbTBC9 is reported to localize to spotted structures at the nuclear periphery in *T. brucei*, which is likely the nuclear pore.^51^ To determine if this protein could rescue the TgTBC9 knockdown, we also expressed it as a YFP fusion and found that it localizes to both the cytoplasm and parasite nucleus (Figure S3C). Although TbTBC9 was expressed at the correct size, plaque assays showed that the TbTBC9 protein could not rescue function of the knockdown at all (Figure S3D-G). This data demonstrates that while the N-terminal YFP tagged TgTBC9 and PfTBC9 can rescue function, another similar RootA orthologue is unable to do so.

### Identification of candidate Rabs that are targeted by TgTBC9

Having demonstrated that the catalytic activity of TgTBC9 is required for function, we next sought to determine which of the Rab proteins that are involved in vesicle trafficking are targeted by TgTBC9. To identify candidate Rabs as well as other potential binding partners, we carried out large scale immunoprecipitations of TgTBC9^3xHA^ via its C-terminal epitope tag. The bound proteins were eluted using high pH and the eluates analyzed by SDS-PAGE, which showed a strong enrichment of the target protein (Figure 6A). The eluted proteins were then identified by mass spectrometry which showed that TgTBC9 was the top hit in the pull down (Figure 6B; Table S3). We also identified Rab1A, Rab2, Rab5A, Rab6 and Rab11B as candidate small GTPase interactors. Of these candidates, Rab2 stood out as the most likely target of TgTBC9 as it was significantly more enriched in the dataset, localizes to the ER/Golgi, and has been shown to be essential for growth using an overexpression screen.^17^ The mass spectrometry dataset also appeared to be enriched in secretory proteins, including a number of rhoptry, dense granule, and IMC proteins, which likely represents cargo trafficking through the secretory pathway that co-immunoprecipitated with TgTBC9 (Table S3).

**Figure 6.**
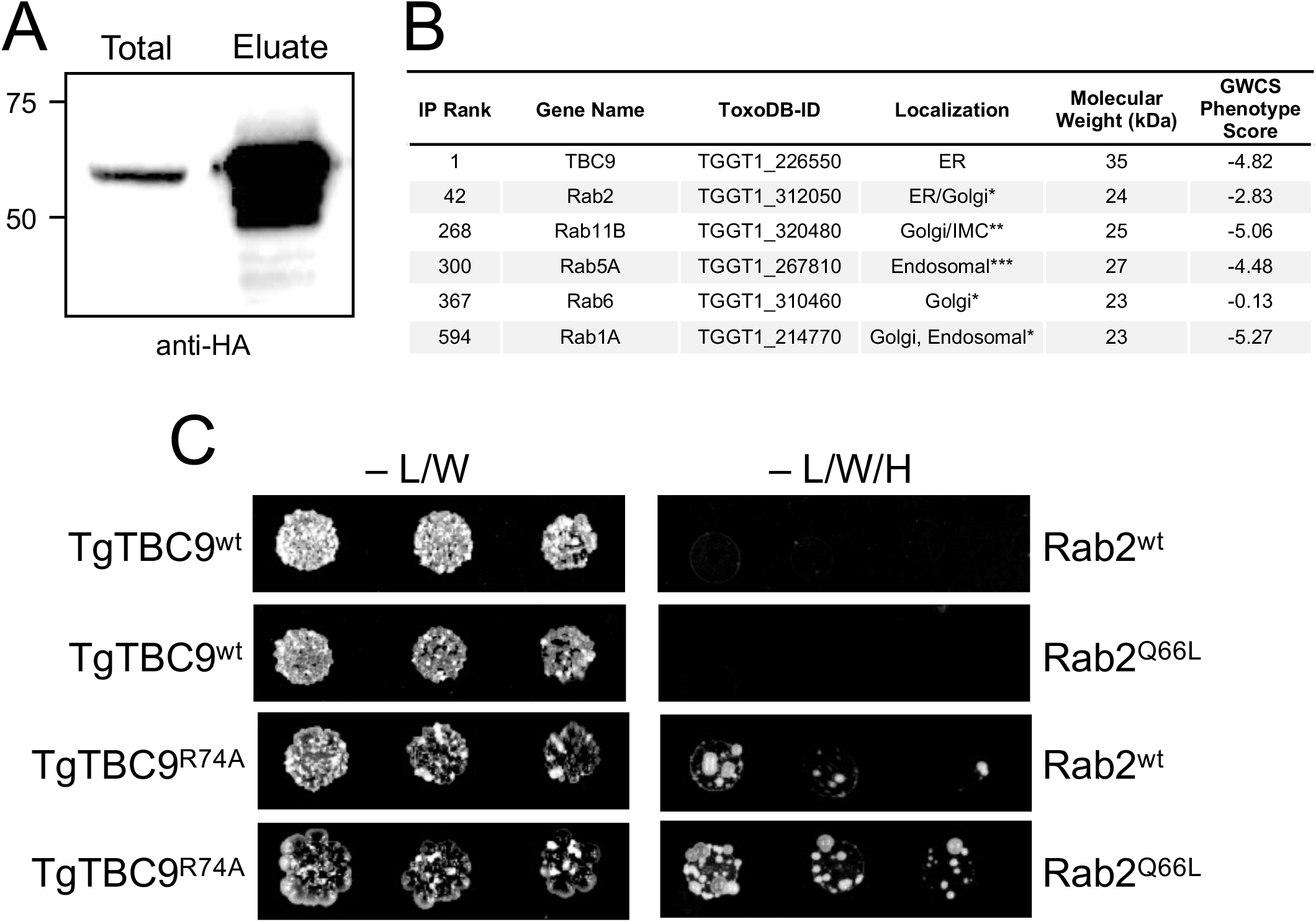
Immunoprecipitation and pairwise Y2H of TgTBC9 reveals Rab2 an interactor. A) Western blot analysis of the TgTBC9 immunoprecipitation showing the input (Total) and eluted material (Eluate) probed with mouse anti-HA antibodies. B) Table showing the rank of TgTBC9 and small GTPase proteins from the IP analysis. The complete dataset is shown in Table S2. Untagged parasites were used as the control. The ToxoDB-IDs, localization (* determined in ^17^, ** determined in ^21^, *** determined in ^20^), molecular weight, and GWCS phenotype score are also shown.^43^ C) Spot assays of pairwise Y2H assessing TgTBC9 and Rab2 interaction using either wild-type (wt), GTP-locked mutant (Rab2^Q66L^), or catalytical inactive mutant (TgTBC9^R74A^) sequences. Yeast expressing the indicated constructs were grown under permissive (−L/W) or restrictive (−L/W/H) conditions to assess interactions.

### TgTBC9 directly interacts with Rab2

To determine if TgTBC9 directly interacts with Rab2, we carried out pairwise yeast two-hybrid (Y2H) experiments. The interactions of TBC proteins with their targets are often transient, thus these interactions are typically assessed using either the catalytically inactive TBC protein, a GTP-locked mutant of the Rab protein, or both mutants.^35^ The rationale is that these mutants enable binding of the GAP with its partner Rab without GTP hydrolysis, leading to an improved interaction by Y2H.^35^ Thus, we used both the wild-type and the catalytically inactive (R74A) mutant of TgTBC9 as bait and either the wild-type or the GTP-locked (Q66L) mutant of Rab2 as prey in the Y2H assay. Using this approach, we found that the TgTBC9^wt^ did not interact with either Rab2^wt^ or Rab2^Q66L^ (Figure 6C). However, the inactive TgTBC9^R74A^ mutant interacts with both Rab2^wt^ or Rab2^Q66L^, with the GTP-locked mutant showing stronger binding (Figure 6C). This data demonstrates that TgTBC9 directly interacts with Rab2 and indicates that this GAP-Rab pair collaborate to regulate secretory traffic in the parasite.

## Discussion

In this study, we explore the localization and function of the *T. gondii* TBC proteins. *Toxoplasma* contains substantially fewer TBC proteins than mammalian cells, which agrees with apicomplexans only containing a reduced set of core Rab proteins.^18,52^ This smaller number of regulators and their targets may simplify characterization of how these proteins regulate protein trafficking throughout the secretory pathway and to the unique organelles of the parasite. Of the 18 predicted TBC proteins in *T. gondii*, 16 are expressed in tachyzoites with various localizations, including the ER, Golgi, cytoplasmic vesicles, and maternal and daughter bud IMC. Those that localize to the ER versus cytoplasmic vesicles are difficult to differentiate as not all proteins reported as ER proteins in *T. gondii* have the classic reticular pattern encircling the nucleus like SERCA.^39,45,53^ Those that localize to cytoplasmic vesicles are likely to partially overlap with different components of the endocytic and exocytic pathways, which intersect downstream of the Golgi.^13,22^ However, some of these could reside in other types of vesicles that are distinct from the endocytic/endocytic pathways as TBC proteins have been shown to function outside of these pathways in other systems.^26,54,55^ More detailed studies of each of these will resolve their precise localization, and they may serve as good markers for further delineating these compartments or subcompartments within the parasite. The utility of TBC proteins markers is likely also true for the Golgi, where the four family members have slightly different localizations indicative of medial, trans, or post-Golgi compartments. Interestingly, the localizations of the TBC proteins in *Toxoplasma* align well with that of the Rabs, which have also been shown to localize to the ER, Golgi, cytoplasmic vesicles, and IMC.^17,20–22^ It is also intriguing that the TBC and Rab proteins are not present in the downstream secretory organelles (i.e. micronemes, rhoptries, or dense granules), suggesting other regulators mediate vesicular traffic in these compartments. Ultimately, determining the precise function of each TBC protein and identifying their target(s) will be important for a complete understanding of secretory and vesicular traffic in *T. gondii* and related parasites.

The fact that the number of TBC proteins exceeds that of the Rabs and that many of the TBC proteins are predicted to be dispensable suggests that there are redundancies among family members.^17,43^ Redundancy would be most likely in regions of the parasite that contain multiple family members such as the Golgi, cytoplasmic vesicles, or daughter IMC, indicating that multiple knockouts may be required to assess function. One likely candidate for redundancy is the TBS proteins TgTBC16 and TgTBC17, which both localize to cytoplasmic vesicles and are each predicted to be dispensable.^32,43^ Another possibility is TgTBC11 and TgTBC15, which localize to IMC daughter buds indicating a role in replication; however, TgTBC11 has a −2.83 phenotype score in the GWCS, suggesting disruption of its gene alone may significantly impair the parasites.^43^ Other proteins with relatively negative phenotype scores include TgTBC18 (−2.86), TgTBC13 (−2.04) and TgTBC2 (−1.84) which may yield significant phenotypes when disrupted individually; however, they also have other family members nearby that may compensate or partially compensate for their absence.^43^ We also cannot exclude the possibility that other regulators lacking TBC domains exist, some TBC proteins carry out other functions, or the intrinsic GTPase activities of some of the Rabs are sufficient for function.^16,27,56^

While many TBC proteins in both *T. gondii* and other systems appear to be dispensable, we demonstrate here that TgTBC9 is essential for growth *in vitro*. TgTBC9 is the smallest member of the family and largely consists of its TBC domain, which we show requires both of its conserved active site residues for function. Conditional knockdown of the protein arrests parasite growth and disrupts the organization of the ER and Golgi-associated compartments, but it does not have a gross effect on the micronemes and rhoptries. The phenotype of the knockdown differs from the knockdown of the sortilin-like receptor protein TgSORTLN which disrupts the trafficking of secretory cargo to the micronemes and rhoptries but does not result in growth arrest.^57^ Similarly, knockdown of the retromer complex in *T. gondii* results in a loss of the parasite’s secretory organelles and mistargeting of their cargo.^58,59^ Although retromer knockdown also results in defects in parasite replication and morphology, it does not cause growth arrest. Disruption of other regulators of secretory traffic such as Rab11A, which affects dense granule and plasma membrane transport, and Rab11B, which disrupts biogenesis of the IMC, also do not arrest growth.^21,22^ The phenotype we observe for TgTBC9 may be a consequence of a disruption of the earliest stages of secretory traffic that results in an ER stress feedback loop which blocks replication as well as the delivery of newly synthesized cargo from entering into the secretory pathway.^60^ Our identification of at least two other ER family members indicates that these TBC proteins cannot compensate for the loss of TgTBC9 and likely carry out distinct functions in the ER.

Phylogenetic analyses of TBC proteins have proven challenging, likely because of ancient divergences among the ancestral clades present in the most recent common ancestor of eukaryotes, which results in homoplasy and problems with alignment hampering phylogenetic inference.^25^ Ancestral TBC clades have experienced differential retention and diversification in different eukaryotic lineages, and as a result the phylogenetic relations of homologs within particular TBC clades is less difficult to infer, typically following the phylogenetic relationships of the taxa of origin. Our phylogeny of TBC-RootA proteins confirms that TgTBC9 belongs to the “RootA” clade of TBC proteins, which was first described in studies examining *T. brucei* TBC proteins and appears to be restricted to protozoans.^25^ The *Plasmodium* orthologue of TgTBC9 is well conserved, predicted to be essential, and can rescue the lethal TgTBC9 knockdown, suggesting this protein has a conserved function in apicomplexans.^50^ This is likely not the case for the *T. brucei* family member which has been experimentally determined to be nuclear in *T. brucei* and cannot rescue the TgTBC9 knockdown.^51^ It will be interesting to determine the localizations and functions of the other RootA family members to determine how this clade has diversified to carry out distinct functions in the protozoa.

We used immunoprecipitation and Y2H analyses to identify Rab2 as a direct binding partner and likely target of TgTBC9. Rab2 is a good fit for a TgTBC9 target as it has been shown to localize to the early secretory pathway (ER, Golgi) and is known to control anterograde traffic between the ER and Golgi.^17,61^ In addition, the overexpression of Rab2 in *T. gondii* indicated that it is essential for growth with no apparent effects the micronemes or rhoptries.^17^ Thus, it appears that Rab2 overexpression essentially phenocopies the knockdown of TgTBC9. Future experiments will focus on biochemical studies on the GTPase activity of TgTBC9 on Rab2 and a more detailed phenotypic analysis of Rab2 using overexpression of dominant active and dominant negative versions of the protein. Since some TBC proteins in other systems are able to act on multiple Rabs, we cannot exclude the possibility that TgTBC9 also regulates other Rabs in the *Toxoplasma* secretory pathway.^62,63^ Other potential targets that were lower ranked in our IP analysis include Rab1A, Rab5A, Rab6 and Rab11B, which localize to the Golgi or endosomal pathways.^17,20,21^ However, there are also multiple TBC proteins in these areas that could regulate these GTPases.

Taken together, this study sheds light on the localization and function of TBC proteins in *T. gondii*. Characterization of the TBC proteins contributes to a broader understanding of the secretary pathway and how proteins are trafficked in the parasite. While redundancy may exist in certain regions of the secretory or vesicular trafficking pathways, we demonstrate that TgTBC9 is essential for growth *in vitro* and plays a crucial role in regulating the early stages of secretory traffic. Our findings also indicate that TgTBC9 directly targets Rab2, a key GTPase involved in the ER-Golgi trafficking pathway. Because TgTBC9 is protozoan-specific and may have activities that are distinct from its mammalian counterparts, it could serve as a new therapeutic target that can be used to combat *T. gondii* and related apicomplexan parasites.

## Acknowledgements

We thank Dr. David Sibley for generously sharing the mouse anti-SERCA antibody, Dr. Gary Ward for sharing the mouse anti-IMC1 antibody, Dr. Gustavo Arrizabalaga for sharing the guinea pig anti-NHE3 antibody, Dr. Kent Hill for *T. brucei* genomic DNA, and Dr. Michael Reese for assistance with the Y2H experiments. We also thank members of the Bradley lab for input on the project as well as reading and commenting on the manuscript. This work was supported by NIH grants AI123360 to P.J.B, and GM089778 to J.A.W, as well as startup funds from UCLA and IU to L.A.N. In addition, J.J.Q was supported by a Milton Gottlieb endowment award and by a Beckman Scholars Award through the Arnold and Mabel Beckman Foundation. The funders had no role in the study design, data collection, interpretation, or decision to submit the work for publication.

## Materials and Methods

### *Toxoplasma* and host cell culture

Parental *T. gondii* RHΔ*ku80*Δ*hpt* and modified strains were grown on confluent monolayers of human foreskin fibroblasts (HFF) host cells at 37°C and 5% CO_2_ in Dulbecco’s modified Eagle medium (DMEM) supplemented with 5% fetal bovine serum (Gibco), 5% Cosmic Calf serum (HyClone), and penicillin-streptomycin-l-glutamine (Gibco). Constructs containing selectable markers were selected using 40 μM chloramphenicol (CAT), 1 μM pyrimethamine, or 50 μg/ml mycophenolic acid-xanthine (HXGPRT).^64–66^ Homologous recombination to the UPRT locus was negatively selected using 5 μM 5-fluorodeoxyuridine (FUDR).^67^

### Sequence analysis and phylogenetic inference

Sequences for TgTBC1-18 from *T. gondii* were retrieved from ToxoDBv43 (https://toxodb.org/toxo/) and were assessed for TBC and other domains by NCBI conserved domain search tool, SMART, and Interpro.^28–30^ Ortholog groups were retrieved from OrthoMCL database (https://orthomcl.org/orthomcl/app) and further assessed by NCBI BLAST analysis. Accession numbers for RootA sequences are given in Table S2 and multiple sequence alignments were generated in Geneious Prime 2021.2.2. (https://www.geneious.com). Maximum Likelihood phylogenetic inference of the ∼200 amino acid TBC-domain of the RootA TBC proteins was preformed in RAxML using the LG+G4 amino acid substitution model, estimating support over 1000 bootstrap replicates, and results were visualized by the Interactive Tree of Life (iTOL) program.^68,69^ Protein alignments were generated using Clustal Omega.^70^

### Chemicals and Antibodies

3-Indoleacetic acid (IAA; Sigma-Aldrich; I2886) was used at 500 µM from a 500 mM stock in 100% ethanol. The hemagglutinin (HA) epitope was detected with mouse monoclonal antibody (mAb) anti-HA (HA.11) (BioLegend) or rabbit mAb anti-HA (C29F4, Cell Signaling). The Ty1 epitope was detected with mouse mAb BB2^71^, V5 epitope was detected with mouse mAb anti-V5 (#R96025; Invitrogen), and the OLLAS tag was detected using the rat mAb anti-OLLAS.^72^ The c-Myc epitope was detected with mouse anti-Myc (mAb 9E10) or rabbit polyclonal antibody (pAb) (#PA1981; Invitrogen). *Toxoplasma*-specific antibodies include rabbit pAb anti-IMC6^73^, mouse anti-SERCA^39^, mouse mAb anti-F1β subunit (5F4)^74^, mouse mAb anti-ATrx1 (11G8)^75^, mouse mAb anti-MIC2^76^, mouse mAb anti-ROP7 (1B10)^77^, mouse pAb anti-Gra14^77^, mouse anti-IMC1^78^, and guinea pig anti-NHE3.^12^

### Immunofluorescence assays and Western blotting

HFF cells were grown to confluency on glass coverslips and infected with *T. gondii* parasites expressing previously mentioned epitope tags. After time periods ranging between 18-36 hours, infected coverslips were fixed using 3.7% formaldehyde and processed for indirect immunofluorescence assay (IFA) as described previously.^79^ Primary antibodies were detected by species-specific secondary antibodies conjugated to Alexa594/488. Coverslips were mounted in Vectashield (Vector Labs) and viewed with an Axio Imager.Z1 fluorescence microscope (Zeiss). Images were processed with the ZEN 2.3 software (Zeiss), which included deconvolution and adaptation of the magenta pseudocolor from the 594 fluorophores.

For Western blotting, parasites were lysed in 1x Laemmli sample buffer (50 mM Tris-HCl [pH 6.8], 10% glycerol, 2% SDS, 100 mM DTT, 0.1% bromophenol blue) and boiled at 100°C for 10 min. Lysates were resolved by SDS-PAGE and transferred to nitrocellulose membranes. Protein blots were probed with the appropriate primary antibody followed by the corresponding secondary antibodies conjugated to horse radish peroxidase (HRP). Proteins were visualized by chemiluminescence (Thermo Scientific) and imaged on ChemiDoc XRS+ (Bio-Rad).

### Epitope tagging

For endogenous C-terminal tagging, guides approximately 100 to 200 bp downstream of each gene’s stop codon were ligated into a pU6-Universal plasmid, as described previously.^80^ A homology-directed repair (HDR) template was amplified from LIC (ligation-independent cloning) epitope-tagging vectors (e.g., 3xHA, smHA, mIAA7-3xHA, 3xMyc, smMyc, 3xV5, 3xTy, and smOLLAS) with 40-bp flanking regions for recombination at the 3′ untranslated region (UTR) of each gene and a selection cassette.^81^ This template was PCR amplified in a 400 μl total reaction, purified by phenol-chloroform extraction, precipitated using ethanol, and electroporated into RHΔ*hxgprt*Δ*ku80* parasites, along with 50 μg of the sequence-verified pU6-Universal plasmid. Confluent monolayers of HFF cells were infected with transfected parasites, and appropriate drug selection (containing either 1 μM pyrimethamine, 50 μg/ml mycophenolic acid/xanthine, or 1 μM chloramphenicol) was applied. Parasites that underwent successful tagging were screened by IFA, and clonal lines of tagged parasites were obtained through limiting dilution. TgTBC1-18 were tagged using CRISPR/Cas9 with primers P1 to P72 and verified by PCR using primers P73 to P92 (Table S4). TgTrs85, TgVps9, Der1-2, TgCep250, and IMC29 were tagged with primers P123 to P142 (Table S4).^11,23,45,82^

### Plaque assays

For plaque assays, six-well plates with HFF monolayers were allowed to reach confluency. Equivalent number of parasites were grown on confluent HFF monolayers ± 500µM IAA. Parasite plaques were allowed to form for 7 days, after which cells were fixed with ice-cold methanol and stained with crystal violet.^83^ The areas of 50 plaques per condition was measured using ZEN software (Zeiss). All plaque assays were performed in triplicate using biological replicates. Statistical significance was calculated using a two-sample two-tailed *t*-test. Graphs and figures were generated using Prism GraphPad 8.0.

### Complementation and mutant construct generation

To generate the TgTBC9 wild-type complement construct, the entire coding region of the gene was PCR amplified from cDNA and cloned into a UPRT-locus knockout vector driven by the TgGT1_238895 promoter, as previously described (primers P93-P96; Table S4).^84^ The online NEBasechanger (https://nebasechanger.neb.com) was used to design the primers. Mutant constructs were built with the complementation vector as a template using the Q5 Site-Directed Mutagenesis Kit (NEB) to mutate specific amino acids (primers P97-P100; Table S4). The plasmids constructs were linearized with PsiI-v2 (NEB), transfected into the TgTBC9^AID^ parasites along with a universal pU6 tagging to the UPRT coding region, and selected with 5µg/ml 5-Fluoro-5’-deoxyuridine (FUDR), as previously described.^67^ TgTBC9^wt-3xTy^, TgTBC9^R74A-3xTy^, or TgTBC9^Q101A-3xTy^ expressing clones were confirmed by IFA.

The TbTBC9 complementation construct was amplified from genomic DNA and cloned into a UPRT-locus knockout vector driven from a tubulin promoter (primers P109-P112; Table S4). The PfTBC9 gene was codon optimized using the Codon Optimization Tool (https://www.idtdna.com/pages/tools/codon-optimization-tool) and synthesized by IDT. The syntenic gene block was cloned into UPRT-locus vectors (primers P101-P108; Table S4).

### Immunoprecipitation

For immunoprecipitation, TgTBC9^3xHA^ or control parasites (RHΔ*hpt*Δ*ku80*) were used to infect ten T150 plates with confluent monolayers of HFF. Extracellular parasites were collected, washed in PBS, and lysed using NP-40 lysis buffer (50 mM Tris [pH 8.0], 150 mM NaCl, 1% NP-40) supplemented with Complete Protease Inhibitor Cocktail (Roche) for 30 minutes on ice. Lysates were centrifuged at 10,000 x *g* for 15 minutes to pellet and remove insoluble material after which the remaining soluble lysate was incubated with anti-HA sepharose beads (Roche) for 4 hours at room temperature. The beads were collected by centrifugation and washed 4 times using NP-40 lysis buffer. Eluted proteins were analyzed by Western blot, and the remaining material was analyzed by mass spectrometry.

### Sample Preparation and LC-MS Acquisition and Analysis

The protein pellets were resuspended with 100ul digestion buffer (8M Urea, 0.1M Tris-HCl pH 8.5), incubated in RT for 30 minutes. Each sample was reduced and alkylated via sequential 20-minute incubations with 5 mM TCEP and 10 mM iodoacetamide at room temperature in the dark while being mixed at 1200 rpm in an Eppendorf thermomixer. 10 μl of carboxylate-modified magnetic beads (CMMB and also widely known as SP3) was added to each sample. ^85^ Ethanol was added to a concentration of 50% to induce protein binding to CMMB. CMMB were washed 3 times with 80% ethanol and then resuspended with 50μl 50mM TEAB.

The protein was digested overnight with 0.1 μg LysC (Promega) and 0.8 μg trypsin (Thermo Scientific, 90057) at 37°C. Following digestion, 1.2ml of 100% acetonitrile was added to each to sample to increase the final acetonitrile concentration to over 95% to induce peptide binding to CMMB. CMMB were then washed 3 times with 100% acetonitrile and the peptide was eluted with 65 μl of 2% DMSO. Eluted peptide samples were dried by vacuum centrifugation and reconstituted in 5% formic acid before analysis by LC-MS/MS.

Peptide samples were separated on a 75μM ID, 25cm C18 column packed with 1.9 μM C18 particles (Dr. Maisch GmbH) using a 140-minute gradient of increasing acetonitrile concentration and injected into a Thermo Orbitrap-Fusion Lumos Tribrid mass spectrometer. MS/MS spectra were acquired using Data Dependent Acquisition (DDA) mode. MS/MS database searching was performed using MaxQuant (1.6.10.43) against the *T. gondii* GT1 reference proteome from ToxoDB.^86^

### Pairwise yeast two-hybrid

Wild-type or the GTP-fixed mutant of Rab2 lacking their C-terminal prenylation sites (residues 1-212) were cloned into the pP6 vector (Hybrigenics SA) as N-terminal fusions with the GAL4 activating domain using Gibson assembly (primers P117-P120; Table S4). TgTBC9^R74A^ or wild-type TgTBC9 were cloned into the pB27 vector (Hybrigenics SA) as N-terminal fusions with the LexA DNA binding domain (primers P113-P116; Table S4). The Rab2 mutant construct were produced by site-directed mutagenesis as described above (primers P121 and P122; Table S4). To test for interactions, pairs of constructs were transformed into the L40 strain of *S. cerevisiae* [MATa his3Δ200trp1-901 leu2-3112 ade2 LYS2::(4lexAop-HIS3) URA3::(8lexAop-lacZ) GAL4]. Strains were grown overnight in permissive (−Leu/−Trp) medium, diluted to an OD_600_ of 2, and spotted in 2x dilutions in both permissive and restrictive (−Leu/−Trp/−His) media. Relative growth in the two conditions was assessed after 6 days incubation at 30°C.

## Supplemental Figure Legends

**Figure S1.**
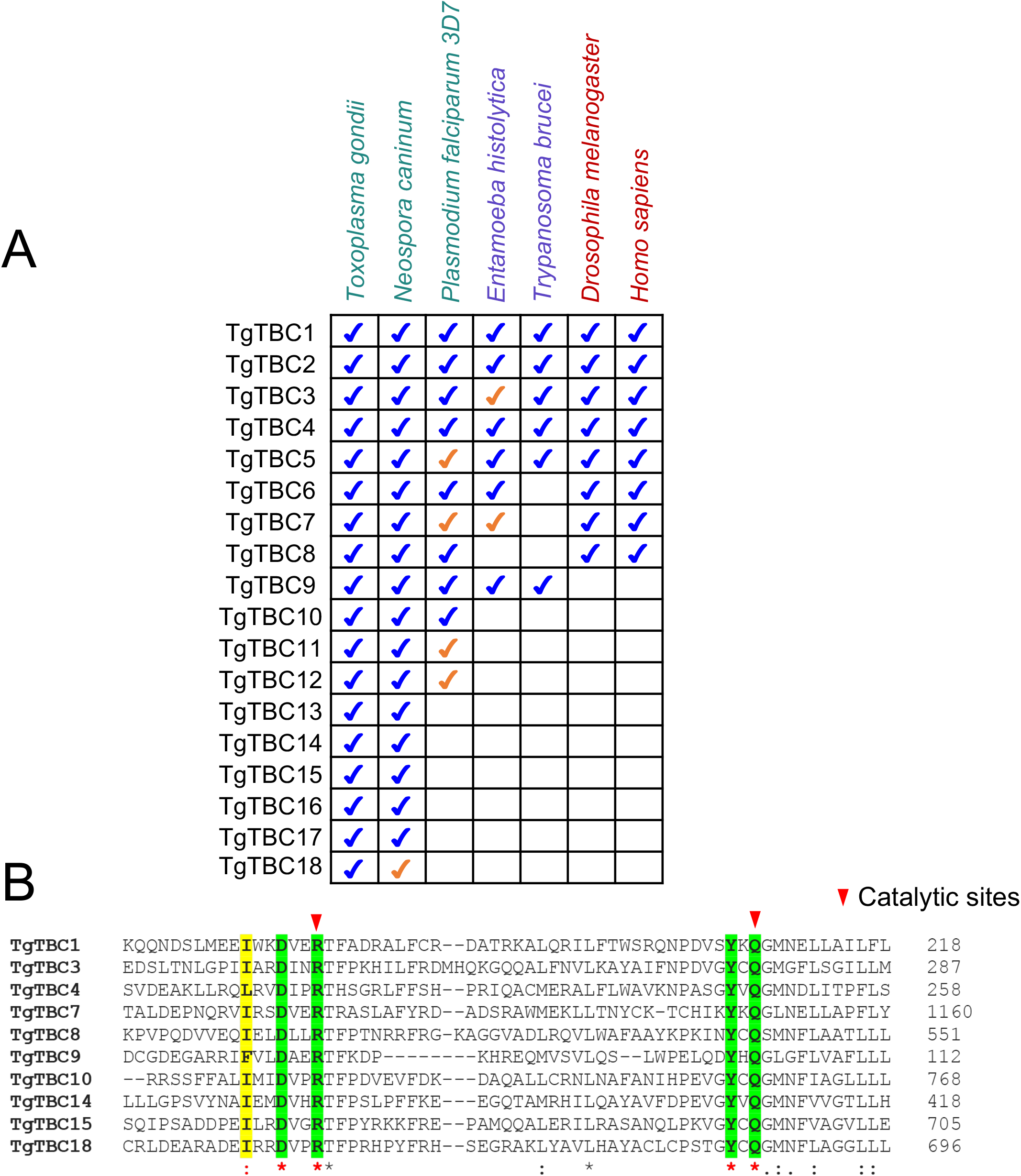
TgTBC1-18 contain orthologs in other systems and most retain the TBC dual-finger active site. A) Diagram of TgTBC1-18 showing orthologs in other species indicated. Blue checks indicate orthologs determined by OrthoMCL, while orange checks indicate likely orthologs based on NCBI BLAST analysis. B) Clustal Omega alignment with the TBC domain region of TgTBC1, 3, 4, 7, 8, 9, 10, 14, 15, and 18 sequences showing TBC dual-finger active site IxxDxxR and YxQ. Bold residues highlighted in yellow depict semi-conserved residues; bold residues in green depict strictly conserved residues. Red triangles indicate the conserved R and Q resides important for catalytic activity. Asterisks (*) indicate identity, colon (:) indicates highly conserved residues, and a period (.) indicates weakly conserved residues.

**Figure S2.**
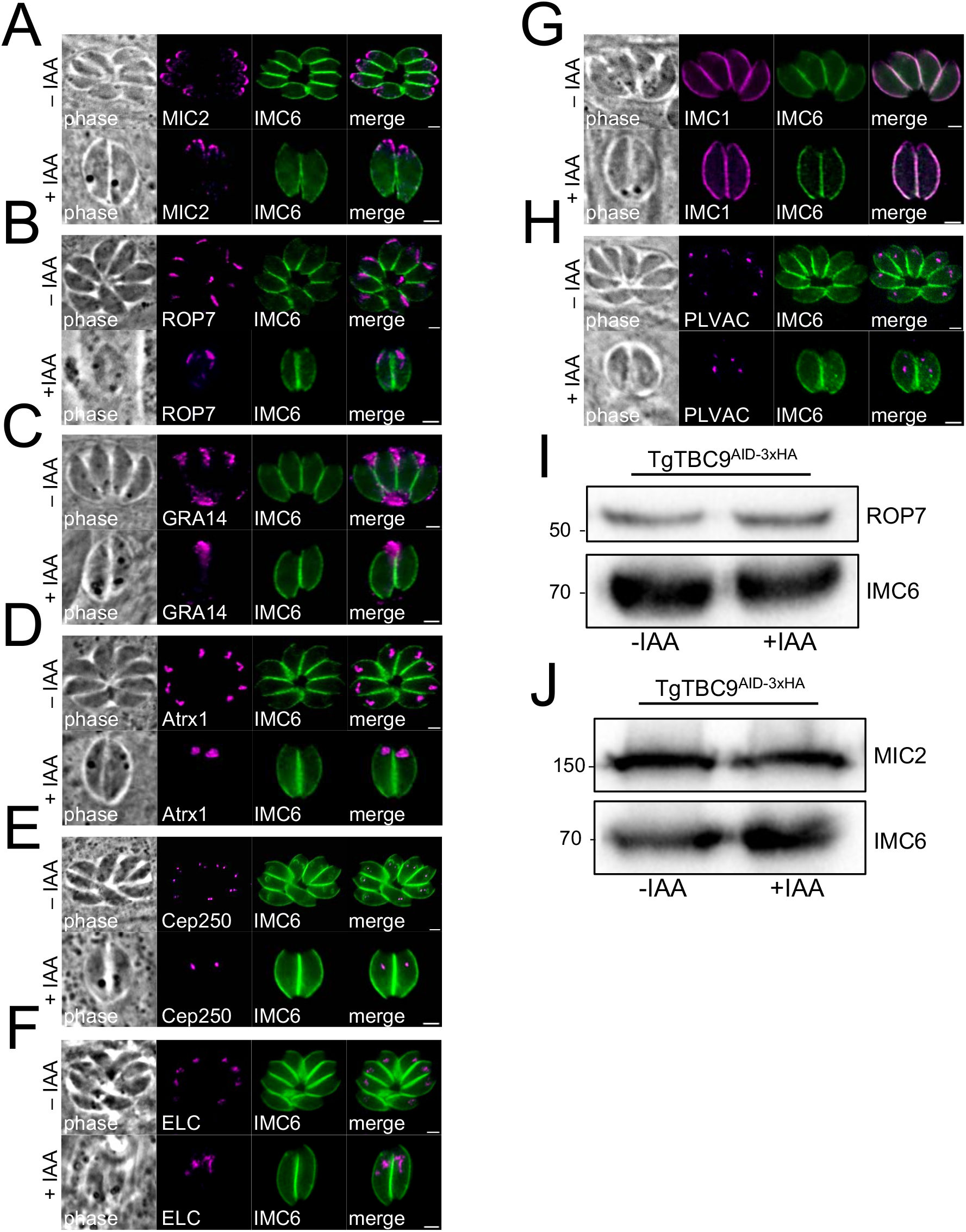
Organelles unaffected by knockdown of TgTBC9. A) IFA of TgTBC9^AID^ without (−) or with (+) IAA for 24 h (following a 4 h pretreatment in ±IAA) showing that the micronemes are unaffected using anti-MIC2. Magenta, mouse anti-MIC2; green, rabbit anti-IMC6. B) IFA of TgTBC9^AID^ parasites grown as described in A but staining for rhoptries using anti-ROP7. Magenta, mouse anti-ROP7; green, rabbit anti-IMC6. C) TgTBC9^AID^ parasites grown as described in A but with staining for dense granules using anti-GRA14. Magenta, mouse anti-GRA14; green, rabbit anti-IMC6. D) TgTBC9^AID^ parasites grown as described in A but with staining for apicoplast using anti-Atrx1. Magenta, mouse anti-Atrx1; green, rabbit anti-IMC6. E) TgTBC9^AID^ parasites grown as described in A but with staining for centrosomes using an endogenously tagged Cep250^3xV5^ strain. Magenta, mouse anti-V5; green, rabbit anti-IMC6. F) TgTBC9^AID^ parasites grown as described in A but with staining for the ELC using an endogenously tagged Vps9^3xV5^ strain. Magenta, mouse anti-V5; green, rabbit anti-IMC6. G) TgTBC9^AID^ parasites grown as described in A but with staining for the IMC. Magenta, mouse anti-IMC1; green, rabbit anti-IMC6. H) TgTBC9^AID^ parasites grown as described in A but with staining for the PLVAC using anti-NHE3. Magenta, guinea pig anti-NHE3; green, rabbit anti-IMC6. Scale bars for all images, 2 μm. I) Western blot analysis of showing ROP7 protein levels are unaffected upon IAA treatment. IMC6 is used as a load control. J) Western blot analysis of showing MIC2 proteins levels upon IAA treatment. IMC6 is used as a load control.

**Figure S3.**
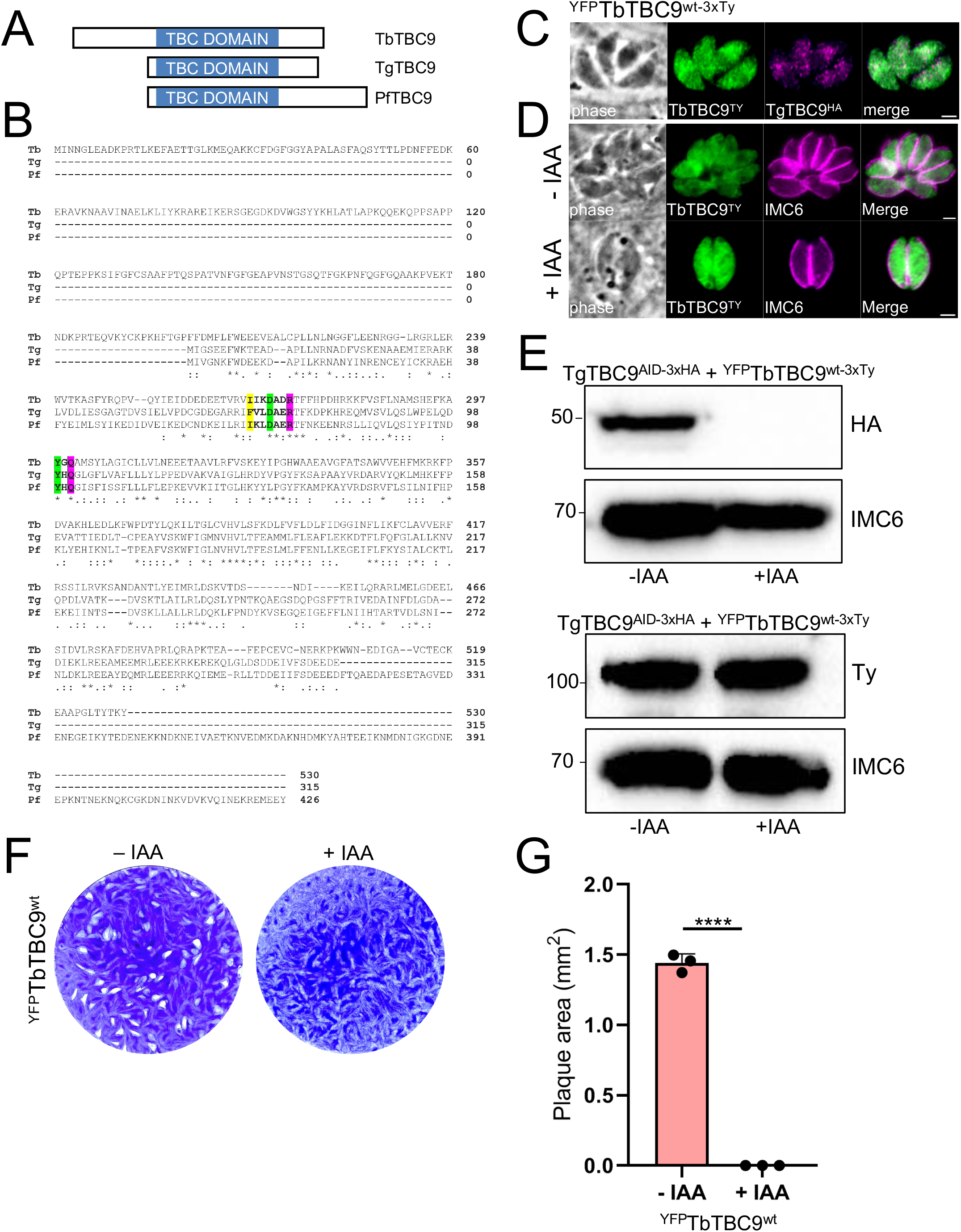
TbTBC9 does not rescue the lethal knockdown of TgTBC9. A) Diagram showing alignment of TbTBC9, TgTBC9, and PfTBC9 revealing N-terminal extension in TbTBC9 and C-terminal extension in PfTBC9. B) Full protein alignment of TbTBC9 (Tb), TgTBC9 (Tg) and PfTBC9 (Pf) using Clustal Omega.^70^ Bold residues highlighted in yellow depict semi-conserved residues; bold residues in green depict strictly conserved residues; bold residues in magenta indicate R and Q residues important for catalytic activity. C) IFA of ^YFP^TbTBC9^wt-3xTy^ colocalized with TgTBC9^AID-3xHA^ showing overlap. Green, mouse anti-Ty; magenta, rabbit anti-HA. Scale bar = 2 μm. D) IFA of TbTBC9^wt^ with (−) or without (+) IAA for 24 h showing that TbTBC9^wt^ is expressed in the presence and absence of IAA. Magenta, mouse anti-Ty; green, rabbit anti-HA. Scale bar = 2 μm. E) Western blot analysis of ^YFP^TbTBC9^wt-3xTy^ in the background of TgTBC9^AID^ tagged parasites. IMC6 is a loading control. F) Plaque assays showing that TbTBC9^wt^ complemented parasites +IAA fail to form plaques. G) Quantification of plaque area at day 7 showing no plaque formation by TbTBC9^wt^ complemented parasites +IAA (****, *P <* 0.0001).

## References

1. Levine, N. D. Progress in Taxonomy of the Apicomplexan Protozoa. J. Protozool. 35, 518– 520 (1988).

2. Davies, A. P. & Chalmers, R. M. Cryptosporidiosis. BMJ 339, b4168–b4168 (2009).

3. Tenter, A. M., Heckeroth, A. R. & Weiss, L. M. Toxoplasma gondii: from animals to humans. Int. J. Parasitol. 30, 1217–1258 (2000).

4. Miller, L. H., Baruch, D. I., Marsh, K. & Doumbo, O. K. The pathogenic basis of malaria. Nature 415, 673–679 (2002).

5. Blader, I. J., Coleman, B. I., Chen, C.-T. & Gubbels, M.-J. Lytic Cycle of Toxoplasma gondii: 15 Years Later. Annu. Rev. Microbiol. 69, 463–485 (2015).

6. Gubbels, M.-J. & Duraisingh, M. T. Evolution of apicomplexan secretory organelles. Int. J. Parasitol. 42, 1071–1081 (2012).

7. Joiner, K. A. & Roos, D. S. Secretory traffic in the eukaryotic parasite Toxoplasma gondii. J. Cell Biol. 157, 557–563 (2002).

8. Hager, K. M., Striepen, B., Tilney, L. G. & Roos, D. S. The nuclear envelope serves as an intermediary between the ER and Golgi complex in the intracellular parasite Toxoplasma gondii. J. Cell Sci. 112, 2631–2638 (1999).

9. Tomavo, S., Slomianny, C., Meissner, M. & Carruthers, V. B. Protein Trafficking through the Endosomal System Prepares Intracellular Parasites for a Home Invasion. PLoS Pathog. 9, e1003629 (2013).

10. Sheiner, L. & Soldati-Favre, D. Protein Trafficking inside Toxoplasma gondii. Traffic 9, 636– 646 (2008).

11. Cao, S., Yang, J., Fu, J., Chen, H. & Jia, H. The Dissection of SNAREs Reveals Key Factors for Vesicular Trafficking to the Endosome-like Compartment and Apicoplast via the Secretory System in Toxoplasma gondii. mBio 12, e01380–21 (2021).

12. Francia, M. E. et al. A Toxoplasma gondii protein with homology to intracellular type Na+/H+ exchangers is important for osmoregulation and invasion. Exp. Cell Res. 317, 1382–1396 (2011).

13. McGovern, O. L., Rivera-Cuevas, Y., Kannan, G., Narwold, A. J. & Carruthers, V. B. Intersection of endocytic and exocytic systems in *Toxoplasma gondii*. Traffic 19, 336–353 (2018).

14. Stenmark, H. Rab GTPases as coordinators of vesicle traffic. Nat. Rev. Mol. Cell Biol. 10, 513–525 (2009).

15. Hutagalung, A. H. & Novick, P. J. Role of Rab GTPases in Membrane Traffic and Cell Physiology. Physiol. Rev. 91, 119–149 (2011).

16. Frasa, M. A. M., Koessmeier, K. T., Ahmadian, M. R. & Braga, V. M. M. Illuminating the functional and structural repertoire of human TBC/RABGAPs. Nat. Rev. Mol. Cell Biol. 13, 67–73 (2012).

17. Kremer, K. et al. An Overexpression Screen of Toxoplasma gondii Rab-GTPases Reveals Distinct Transport Routes to the Micronemes. PLoS Pathog. 9, e1003213 (2013).

18. Langsley, G. et al. Comparative genomics of the Rab protein family in Apicomplexan parasites. Microbes Infect. 10, 462–470 (2008).

19. Miranda, K. et al. Characterization of a novel organelle in Toxoplasma gondii with similar composition and function to the plant vacuole: Toxoplasma gondii plant-like vacuole. Mol. Microbiol. 76, 1358–1375 (2010).

20. Robibaro, B. et al. Toxoplasma gondii Rab5 enhances cholesterol acquisition from host cells. Cell. Microbiol. 4, 139–152 (2002).

21. Agop-Nersesian, C. et al. Biogenesis of the Inner Membrane Complex Is Dependent on Vesicular Transport by the Alveolate Specific GTPase Rab11B. PLoS Pathog. 6, e1001029 (2010).

22. Venugopal, K. et al. Rab11A regulates dense granule transport and secretion during Toxoplasma gondii invasion of host cells and parasite replication. PLOS Pathog. 16, e1008106 (2020).

23. Sakura, T. et al. A Critical Role for Toxoplasma gondii Vacuolar Protein Sorting VPS9 in Secretory Organelle Biogenesis and Host Infection. Sci. Rep. 6, 38842 (2016).

24. Morlon-Guyot, J., Pastore, S., Berry, L., Lebrun, M. & Daher, W. *Toxoplasma gondii* Vps11, a subunit of HOPS and CORVET tethering complexes, is essential for the biogenesis of secretory organelles. Cell. Microbiol. 17, 1157–1178 (2015).

25. Gabernet-Castello, C., O’Reilly, A. J., Dacks, J. B. & Field, M. C. Evolution of Tre-2/Bub2/Cdc16 (TBC) Rab GTPase-activating proteins. Mol. Biol. Cell 24, 1574–1583 (2013).

26. Fukuda, M. TBC proteins: GAPs for mammalian small GTPase Rab? Biosci. Rep. 31, 159– 168 (2011).

27. Nagano, F. et al. Molecular Cloning and Characterization of the Noncatalytic Subunit of the Rab3 Subfamily-specific GTPase-activating Protein. J. Biol. Chem. 273, 24781–24785 (1998).

28. Marchler-Bauer, A. & Bryant, S. H. CD-Search: protein domain annotations on the fly. Nucleic Acids Res. 32, W327–W331 (2004).

29. El-Gebali, S. et al. The Pfam protein families database in 2019. Nucleic Acids Res. 47, D427–D432 (2019).

30. Letunic, I., Doerks, T. & Bork, P. SMART: recent updates, new developments and status in 2015. Nucleic Acids Res. 43, D257–D260 (2015).

31. Mossessova, E., Gulbis, J. M. & Goldberg, J. Structure of the Guanine Nucleotide Exchange Factor Sec7 Domain of Human Arno and Analysis of the Interaction with ARF GTPase. Cell 92, 415–423 (1998).

32. Mouratou, B. et al. The domain architecture of large guanine nucleotide exchange factors for the small GTP-binding protein Arf. BMC Genomics 6, 20 (2005).

33. Li, L., Stoeckert, C. J. & Roos, D. S. OrthoMCL: Identification of Ortholog Groups for Eukaryotic Genomes. Genome Res. 13, 2178–2189 (2003).

34. Altschul, S. F., Gish, W., Miller, W., Myers, E. W. & Lipman, D. J. Basic local alignment search tool. J. Mol. Biol. 215, 403–410 (1990).

35. TBC-domain GAPs for Rab GTPases accelerate GTP hydrolysis by a dual-finger mechanism | Nature. https://www.nature.com/articles/nature04847.

36. Hortua Triana, M. A., et al. Tagging of Weakly Expressed *Toxoplasma gondii* Calcium-Related Genes with High-Affinity Tags. J. Eukaryot. Microbiol. 65, 709–721 (2018).

37. Huynh, M.-H. & Carruthers, V. B. Tagging of Endogenous Genes in a *Toxoplasma gondii* Strain Lacking Ku80. Eukaryot. Cell 8, 530–539 (2009).

38. Pelletier, L. et al. Golgi biogenesis in Toxoplasma gondii. Nature 418, 548–552 (2002).

39. Nagamune, K., Beatty, W. L. & Sibley, L. D. Artemisinin Induces Calcium-Dependent Protein Secretion in the Protozoan Parasite *Toxoplasma gondii*. Eukaryot. Cell 6, 2147– 2156 (2007).

40. Back, P. S. et al. IMC29 Plays an Important Role in *Toxoplasma* Endodyogeny and Reveals New Components of the Daughter-Enriched IMC Proteome. mBio 14, e03042–22 (2023).

41. Behnke, M. S. et al. Coordinated Progression through Two Subtranscriptomes Underlies the Tachyzoite Cycle of Toxoplasma gondii. PLoS ONE 5, e12354 (2010).

42. Gajria, B. et al. ToxoDB: an integrated Toxoplasma gondii database resource. Nucleic Acids Res. 36, D553–D556 (2007).

43. Sidik, S. M. et al. A Genome-wide CRISPR Screen in Toxoplasma Identifies Essential Apicomplexan Genes. Cell 166, 1423–1435.e12 (2016).

44. Li, S., Prasanna, X., Salo, V. T., Vattulainen, I. & Ikonen, E. An efficient auxin-inducible degron system with low basal degradation in human cells. Nat. Methods 16, 866–869 (2019).

45. Agrawal, S., van Dooren, G. G., Beatty, W. L. & Striepen, B. Genetic Evidence that an Endosymbiont-derived Endoplasmic Reticulum-associated Protein Degradation (ERAD) System Functions in Import of Apicoplast Proteins. J. Biol. Chem. 284, 33683–33691 (2009).

46. Breinich, M. S. et al. A Dynamin Is Required for the Biogenesis of Secretory Organelles in Toxoplasma gondii. Curr. Biol. 19, 277–286 (2009).

47. Mallo, N. et al. Depletion of a *Toxoplasma* porin leads to defects in mitochondrial morphology and contacts with the endoplasmic reticulum. J. Cell Sci. 134, jcs255299 (2021).

48. Jacobs, K., Charvat, R. & Arrizabalaga, G. Identification of Fis1 Interactors in Toxoplasma gondii Reveals a Novel Protein Required for Peripheral Distribution of the Mitochondrion. mBio 11, e02732–19 (2020).

49. Gavriljuk, K. et al. Catalytic mechanism of a mammalian Rab·RabGAP complex in atomic detail. Proc. Natl. Acad. Sci. 109, 21348–21353 (2012).

50. Zhang, M. et al. Uncovering the essential genes of the human malaria parasite *Plasmodium falciparum* by saturation mutagenesis. Science 360, eaap7847 (2018).

51. Goos, C., Dejung, M., Janzen, C. J., Butter, F. & Kramer, S. The nuclear proteome of Trypanosoma brucei. PLOS ONE 12, e0181884 (2017).

52. Quevillon, E. et al. The Plasmodium falciparum family of Rab GTPases. Gene 306, 13–25 (2003).

53. Márquez-Nogueras, K. M., Hortua Triana, M. A., Chasen, N. M., Kuo, I. Y. & Moreno, S. N. Calcium signaling through a transient receptor channel is important for Toxoplasma gondii growth. eLife 10, e63417 (2021).

54. Yonekura, H. et al. Antisense Display: A New Method for Functional Gene Screen and Its Application to Angiogenesis-Related Gene Isolation. Ann. N. Y. Acad. Sci. 947, 382–386 (2006).

55. Nakamura, T. et al. A novel transcriptional unit of the tre oncogene widely expressed in human cancer cells. Oncogene 7, 733–741 (1992).

56. Barr, F. & Lambright, D. G. Rab GEFs and GAPs. Curr. Opin. Cell Biol. 22, 461–470 (2010).

57. Sloves, P.-J. et al. Toxoplasma Sortilin-like Receptor Regulates Protein Transport and Is Essential for Apical Secretory Organelle Biogenesis and Host Infection. Cell Host Microbe 11, 515–527 (2012).

58. Sangaré, L. O. et al. Unconventional endosome-like compartment and retromer complex in Toxoplasma gondii govern parasite integrity and host infection. Nat. Commun. 7, 11191 (2016).

59. McGovern, O. L. & Carruthers, V. B. Toxoplasma Retromer Is Here to Stay. Trends Parasitol. 32, 758–760 (2016).

60. Sano, R. & Reed, J. C. ER stress-induced cell death mechanisms. Biochim. Biophys. Acta BBA - Mol. Cell Res. 1833, 3460–3470 (2013).

61. Tisdale, E. J. & Balch, W. E. Rab2 Is Essential for the Maturation of Pre-Golgi Intermediates. J. Biol. Chem. 271, 29372–29379 (1996).

62. Itoh, T., Satoh, M., Kanno, E. & Fukuda, M. Screening for target Rabs of TBC (Tre-2/Bub2/Cdc16) domain-containing proteins based on their Rab-binding activity. Genes Cells 11, 1023–1037 (2006).

63. Zhang, X.-M., Walsh, B., Mitchell, C. A. & Rowe, T. TBC domain family, member 15 is a novel mammalian Rab GTPase-activating protein with substrate preference for Rab7. Biochem. Biophys. Res. Commun. 335, 154–161 (2005).

64. Donald, R. G. K., Carter, D., Ullman, B. & Roos, D. S. Insertional Tagging, Cloning, and Expression of the Hypoxanthine-Xanthine-Guanine Phosphoribosyltransferase Gene. J. Biol. Chem. 271, 14010–14019 (1996).

65. Kim, K., Soldati, D. & Boothroyd, J. C. Gene Replacement in *Toxoplasma gondii* with Chloramphenicol Acetyltransferase as Selectable Marker. Science 262, 911–914 (1993).

66. Donald, R. G. & Roos, D. S. Stable molecular transformation of Toxoplasma gondii: a selectable dihydrofolate reductase-thymidylate synthase marker based on drug-resistance mutations in malaria. Proc. Natl. Acad. Sci. 90, 11703–11707 (1993).

67. Donald, R. G. & Roos, D. S. Insertional mutagenesis and marker rescue in a protozoan parasite: cloning of the uracil phosphoribosyltransferase locus from Toxoplasma gondii. Proc. Natl. Acad. Sci. 92, 5749–5753 (1995).

68. Letunic, I. & Bork, P. Interactive Tree Of Life (iTOL) v4: recent updates and new developments. Nucleic Acids Res. 47, W256–W259 (2019).

69. Stamatakis, A. RAxML version 8: a tool for phylogenetic analysis and post-analysis of large phylogenies. Bioinformatics 30, 1312–1313 (2014).

70. Sievers, F. et al. Fast, scalable generation of high-quality protein multiple sequence alignments using Clustal Omega. Mol. Syst. Biol. 7, 539 (2011).

71. Bastin, P., Bagherzadeh, A., Matthews, K. R. & Gull, K. A novel epitope tag system to study protein targeting and organelle biogenesis in Trypanosoma brucei. Mol. Biochem. Parasitol. 77, 235–239 (1996).

72. Viswanathan, S. et al. High-performance probes for light and electron microscopy. Nat. Methods 12, 568–576 (2015).

73. Choi, C. P. et al. A photoactivatable crosslinking system reveals protein interactions in the Toxoplasma gondii inner membrane complex. PLoS Biol. 17, e3000475 (2019).

74. Frénal, K., Marq, J.-B., Jacot, D., Polonais, V. & Soldati-Favre, D. Plasticity between MyoC- and MyoA-glideosomes: an example of functional compensation in Toxoplasma gondii invasion. PLoS Pathog. 10, e1004504 (2014).

75. DeRocher, A. E. et al. A thioredoxin family protein of the apicoplast periphery identifies abundant candidate transport vesicles in Toxoplasma gondii. Eukaryot. Cell 7, 1518–1529 (2008).

76. Huynh, M.-H. & Carruthers, V. B. A Toxoplasma gondii Ortholog of *Plasmodium* GAMA Contributes to Parasite Attachment and Cell Invasion. mSphere 1, e00012–16 (2016).

77. Rome, M. E., Beck, J. R., Turetzky, J. M., Webster, P. & Bradley, P. J. Intervacuolar transport and unique topology of GRA14, a novel dense granule protein in Toxoplasma gondii. Infect. Immun. 76, 4865–4875 (2008).

78. Wichroski, M. J., Melton, J. A., Donahue, C. G., Tweten, R. K. & Ward, G. E. *Clostridium septicum* Alpha-Toxin Is Active against the Parasitic Protozoan *Toxoplasma gondii* and Targets Members of the SAG Family of Glycosylphosphatidylinositol-Anchored Surface Proteins. Infect. Immun. 70, 4353–4361 (2002).

79. Bradley, P. J. et al. Proteomic Analysis of Rhoptry Organelles Reveals Many Novel Constituents for Host-Parasite Interactions in Toxoplasma gondii. J. Biol. Chem. 280, 34245–34258 (2005).

80. Nadipuram, S. M. et al. In Vivo Biotinylation of the Toxoplasma Parasitophorous Vacuole Reveals Novel Dense Granule Proteins Important for Parasite Growth and Pathogenesis. mBio 7, (2016).

81. Sidik, S. M., Hackett, C. G., Tran, F., Westwood, N. J. & Lourido, S. Efficient genome engineering of Toxoplasma gondii using CRISPR/Cas9. PloS One 9, e100450 (2014).

82. Chen, C.-T. & Gubbels, M.-J. TgCep250 is dynamically processed through the division cycle and is essential for structural integrity of the *Toxoplasma* centrosome. Mol. Biol. Cell 30, 1160–1169 (2019).

83. Nadipuram, S. M., Thind, A. C., Rayatpisheh, S., Wohlschlegel, J. A. & Bradley, P. J. Proximity biotinylation reveals novel secreted dense granule proteins of Toxoplasma gondii bradyzoites. PloS One 15, e0232552 (2020).

84. Back, P. S., et al. Ancient MAPK ERK7 is regulated by an unusual inhibitory scaffold required for *Toxoplasma* apical complex biogenesis. Proc. Natl. Acad. Sci. 117, 12164–12173 (2020).

85. Hughes, C. S. et al. Ultrasensitive proteome analysis using paramagnetic bead technology. Mol. Syst. Biol. 10, 757 (2014).

86. Cox, J. & Mann, M. MaxQuant enables high peptide identification rates, individualized p.p.b.-range mass accuracies and proteome-wide protein quantification. Nat. Biotechnol. 26, 1367–1372 (2008).

